# Vaccinia virus induces EMT-like transformation and RhoA-mediated mesenchymal migration

**DOI:** 10.1101/2023.02.04.527154

**Authors:** Wei Liu, Jia-Yin Lu, Ya-Jun Wang, Xin-Xin Xu, Yu-Chen Chen, Sai-Xi Yu, Xiao-Wei Xiang, Xue-Zhu Chen, Yaming Jiu, Hai Gao, Mengyao Sheng, Zheng-Jun Chen, Xinyao Hu, Dong Li, Paolo Maiuri, Xinxin Huang, Tianlei Ying, Guo-Liang Xu, Dai-Wen Pang, Zhi-Ling Zhang, Baohong Liu, Yan-Jun Liu

## Abstract

The emerging outbreak of monkeypox is closely associated with the viral infection and spreading, threatening global public health. Virus-induced cell migration facilitates viral transmission. However, high-resolution dynamics and mechanisms underlying this type of cell migration remain unclear. Here, we investigate the motility of cells infected by vaccinia virus (VACV), a close relative of monkeypox, through combining multi-omics analyses and high-resolution live-cell imaging. We find that, upon VACV infection, the epithelial cells undergo EMT-like transformation, during which they lose intercellular junctions and acquire the migratory capacity to promote viral spreading. After transformation, VACV-induced mesenchymal migration is highly dependent on the actin cytoskeleton and RhoA signaling, which is responsible for the depolymerization of robust actin stress fibers, the leading-edge protrusion formation, and the rear-edge recontraction. Our study reveals how poxviruses alter the epithelial phenotype and regulate RhoA signaling to induce fast migration, providing a unique perspective to understand the pathogenesis of poxviruses.

## Introduction

Cell migration, as a fundamental process, is vital for embryonic development, immune cell response, wound healing, and tumor metastasis^1–3^. As a recognized mode of cell migration, mesenchymal migration is characterized by cell polarization to form a leading-edge extension that ensures adhesive interaction with the extracellular matrix, followed by rear-edge retraction and cell body translocation^4,5^. Epithelial-mesenchymal transition (EMT) is a biological process in which polarized epithelial cells lose tight junctions and become invasive mesenchymal phenotype, crucial in blastocyst migration, gastrulation, tissue repair, and carcinogenesis^6^. The significant hallmark of EMT is the loss of E-cadherin directly or indirectly regulated by transcription factors of the SNAIL family, ZEB family, and TWIST family^7^. Many biological processes are engaged in EMT to entitle migratory cell properties, including activation of transcription factors, expressions of specific cell-adhesion molecules, and cytoskeletal rearrangement^6^. Microbial and viral infection can also induce the EMT transformation involved in the etiology of cancer^8,9^. More importantly, virus-induced aberrant cell motility, followed by EMT, can be an efficient way to spread the nascent virus^10–18^, which requires more scholarly attention.

Vaccinia virus (VACV), a large dsDNA virus^19^, can hijack signaling pathways and regulate the actin cytoskeleton in the host, inducing actin-based virus motility and cell motility for progeny virions spreading^10,13,20–22^. VACV-induced actin tails, a way of fast actin polymerization, require activating localized outside-in signaling. Viral B5R-activated Src phosphorylates viral A36R protein, which is essential for recruitment of N-WASP and Arp2/3 complex^22–24^. Subsequently, actin nucleation and polymerization are initial to help virus cell-to-cell spread. VACV-induced cell locomotion, enhanced by TGF-β-independent smad4 signaling^20^, is another remarkable viral cytopathic effect (CPE). Early and late vaccinia gene expression is essential for VACV to manipulate cell migration^10^. Directly binding to RhoA, viral F11L protein blocks RhoA/Rho-associated kinase (ROCK) signaling, resulting in the depolymerization of actin stress fibers and promoting VACV-induced cell migration (VICM)^13^. PDZ-like domain in F11 protein can also inhibit RhoA signaling by interacting with Rho GTPase-activating protein (GAP) Myosin-9A^25^. Loss of F11 expression fails to induce cell motility and reduces the spread of the virus *in vitro* and *vivo*^11,13^. Furthermore, vaccinia growth factor (VGF), a secretory viral protein, enhances rapid and directed cell migration by stimulating EGFR signaling cascade^12^. Similarly, latent membrane protein 1 (LMP1) of Epstein-Barr virus (EBV) activates the ERK-MAPK pathway to induce cell locomotion^17^. However, cell migration is a complicated process involving spatial-temporal cooperation of signal pathways and cytoskeleton^26,27^. How VACV-hijacked signaling coordinates the host cytoskeleton to initiate cell migration remains elusive.

Here, we systematically investigated the dynamic process of VACV-induced cell behavior by combining multi-omics and high-resolution live-cell imaging. We found that VACV-infected cells underwent a phenotypic transformation process and initiated fast cell migration to spread progeny virions efficiently. We also revealed that VACV-regulated actin cytoskeleton through RhoA/ROCK signaling was involved in the whole process of VICM, responsible for depolymerization of robust actin stress fibers, leading-edge protrusion formation, and rear-edge contraction. Collectively, we propose a unique and efficient way of virus spreading by inducing cell migration. Mechanistically, VACV-hijacked RhoA signaling regulates fast cell migration. Our finding sheds light on antiviral drug development and provides potential targets for virus infection prevention and treatment.

## Results

### Vaccinia virus induces EMT-like transformation and cell migration

Viruses need to hijack cellular machinery for themselves during their entire life cycles^12,18^. In these processes, the virus-infected cell would undergo a phenotypic transformation serving as the viral sanctuary. The VACV-infected cell expressing LifeAct-mRuby exhibited a noticeable change in position during one hour of observation, indicating that vaccinia induced fast cell migration compared to the non-infected control cell (Fig. 1a). To conveniently distinguish between infected and non-infected cells, we performed the infection assay with VACV expressing EGFP. And we found that the morphology of VACV-infected cells altered and cell area decreased, suggesting that VACV infection may alter the cell phenotype (Fig. 1b, c). To dissect the virus-induced cell transition, we performed transcriptomes and proteomes analysis of Vero cells after infection. Principal-component analysis (PCA) showed distinct transcriptomes and proteomes in non-infected and VACV-infected cells (Extended Data Fig.1a, b and 2a, b). The 6724 significantly differentially expressed genes (2472 genes upregulated; 4252 genes downregulated) were identified upon VACV infection (Extended Data Fig. 1c). Genes associated with tight junctions were downregulated, and the expression of transcription factors associated with EMT was upregulated in VACV-infected cells, which hinted at the initiation of EMT (Fig. 1d). Precisely, the integrity of the tight junction protein ZO-1 was lost in virus-infected cells (Extended Data Fig. 3a). Furthermore, the enriched Gene Ontology (GO) terms, launched by differentially expressed genes or proteins, including “positive regulation of epithelial to mesenchymal transition” and “regulation of cytoskeleton organization” which directly and indirectly corresponded to the EMT process, were overrepresented in VACV-infected cells (Fig. 1e, Extended Data Fig. 1d-f and 2c-f). Indeed, the expression of some EMT markers, including E-cadherin, N-cadherin, keratin, and ZO-1, was downregulated, and vimentin was upregulated in VACV-infected cells (Fig. 1f, Extended Data Fig. 3c, d). These results demonstrated that the EMT-like process occurred in VACV-infected epithelial cells as these cells lost their cell-cell adhesion and acquired the ability to migrate.

**Figure 1.**
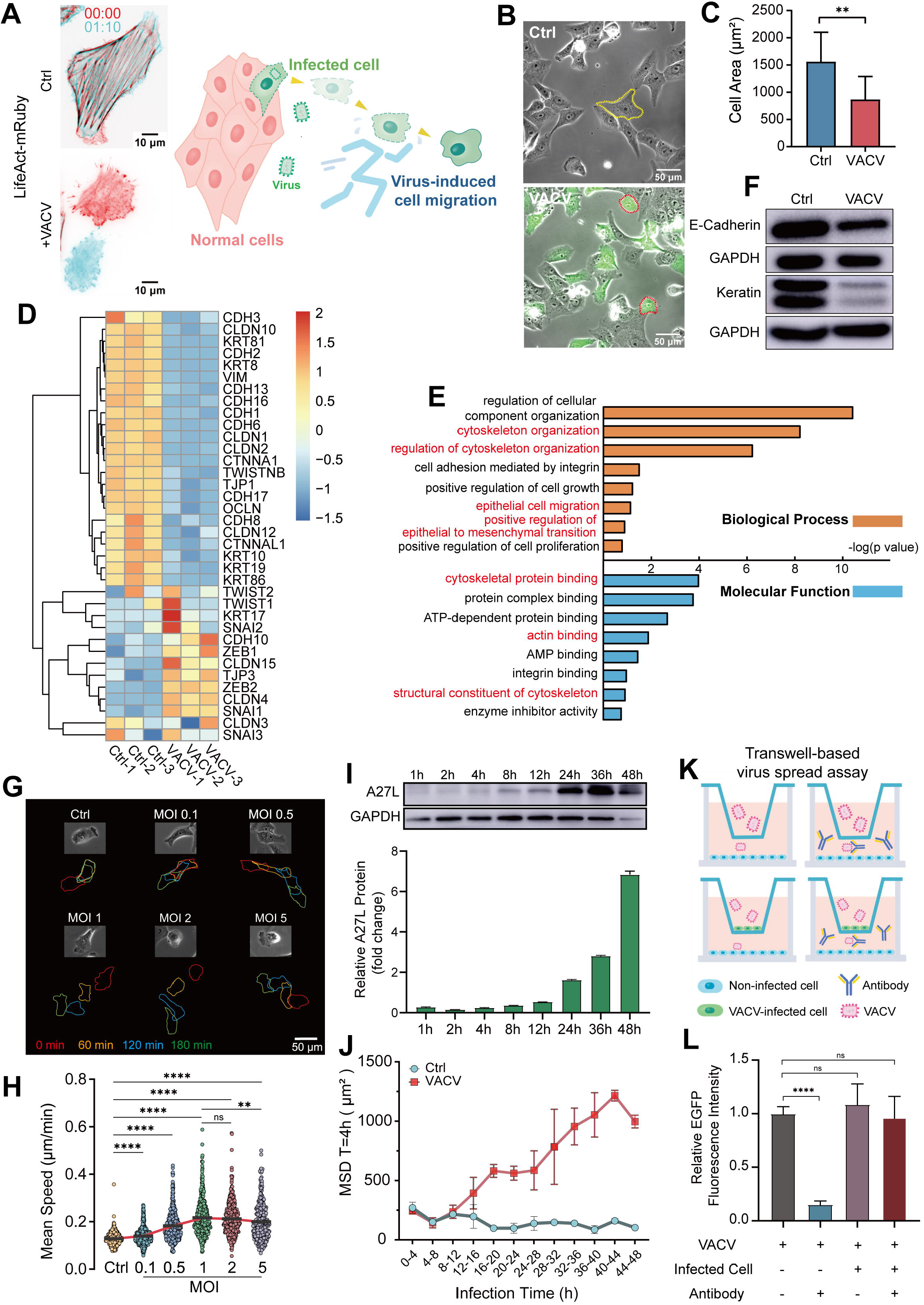
Vaccinia virus induces EMT-like transformation and cell migration. **a**, Representative fluorescent images of a control Vero cell and a VACV-infected Vero cell expressing LifeAct-mRuby (left). Images in red are at t=0 min and in cyan at t= 70 min. Schematic diagram (right) of vaccinia virus-induced cell migration. **b**, Representative phase-contrast images of non-infected Vero cells or VACV-infected Vero cells expressing EGFP. The dotted lines represent the cell outlines (yellow: control cells; red: VACV-infected cells). **c**, Analysis of cell area of control and VACV-infected Vero cells. **d**, Heatmap of genes associated with EMT in control and VACV-infected Vero cells. Each column represents one biological replicate. **e**, Overrepresented GO terms of VACV-infected Vero cells compared with non-infected Vero cells. Highlighted (red) GO terms are associated with EMT and cytoskeletal dynamics. **f**, Immunoblotting of E-cadherin and Keratin in control and VACV-infected cells (n=3). **g**, Representative images of control and VACV-infected cells (different MOI) and corresponding cell outlines at different time points. **h**, Mean speed of control and VACV-infected cells (different MOI). **i**, Immunoblotting (top) and quantitative analysis (bottom) of viral A27L protein during different h.p.i. in Vero cells (n=3). **j**, MSD of moving control (blue) and VACV-infected (red) cells in different periods. **k**, Schematic diagram of transwell-based virus spread assay in vitro. **l**, The fluorescent quantitative analysis of VACV-infected Vero cells in lower chamber at different virus spread conditions. Data are mean ± SEM. *p<0.05, **p<0.01, ***p<0.001, ****p<0.0001 (unpaired t-test).

To optimize the multiplicity of infection (MOI) for VICM, the trajectories of individual cells with different MOI were tracked and analyzed in terms of mean squared displacement (MSD), diffusion coefficient, and mean speed (Fig. 1g, h, Extended Data Fig. 3b, e-g). The curve of mean speed gradually rose to a peak at MOI 1 and did not increase significantly at MOI 2.5 or 5 (Fig. 1h), as well as MSD and diffusion coefficient (Extended Data Fig. 3f, g), which suggests that MOI 1 of VACV was the optimal infective dose of inducing EMT-like process and apparent cell migration. VICM generally appeared at 8 hours post infection (h.p.i). To determine the association between VICM and the viral life cycle, we monitored the expression of A27L (a critical viral protein for the formation of intracellular enveloped virus (IEV) and virus invasion) and the MSD of migrating cells at a fixed time interval (4h) after infection. We found that the A27L gene was expressed, and meanwhile, VICM occurred at approximately 12 h.p.i when the nascent viruses began to assemble (Fig. 1i, j). Hence, we speculated that VICM could be a latent way of spreading the nascent virus. To evaluate virus transmission efficiency with VICM, we performed the transwell-based virus spread assay and found that VACV dissemination was more efficient when VICM occurred (Fig. 1k, l, Extended Data Fig. 3h). More importantly, VACV-neutralizing antibodies had a lower preventive effect against intercellular viral spreading (Fig. 1l). Therefore, virus-infected cells underwent EMT-like transformation and acquired the migratory capacity to spread the nascent virus efficiently.

### VICM is actin-dependent

Cell migration requires the rearrangement of the cytoskeleton, including actin microfilaments, microtubules, and intermediate filament^26^. Transcriptomic and proteomic analysis revealed that VACV infection changed cytoskeleton organization (Fig. 1e, Extended Data Fig. 1f and 2d). To further dissect the cytoskeleton dynamics during VICM, we visualized and analyzed the location and distribution of actin, tubulin, and vimentin. We observed robust actin stress fibers were lost in VACV-infected cells, accompanied by a decrease in actin fluorescence intensity (Fig. 2a, Extended Data Fig. 4a, b). Further fluorescence quantitative analysis along stress fibers demonstrated that the depolymerization of robust actin stress fibers was related to its width reduction (Fig. 2b). These results indicated that depolymerization of robust actin bundles was a prerequisite for VICM. Besides, VACV infection also resulted in abnormal distribution of microtubule and vimentin network (Extended Data Fig. 4a, c). Cells spreading arbitrarily on the glass bottom may lead to various cytoskeleton distributions. Thus, we plated the cells on crossbow-shaped fibronectin micropatterns for uniform spreading, and immunofluorescence analysis further confirmed the loss of strong actin stress fibers (Extended Data Fig. 4d). Myosin, the crucial actin filaments binding protein, plays an essential role in contractile force generation during cell migration^28^. We next tested the expression and distribution of myosin in VACV-infected cells. Although the protein level of myosin light chain (MLC) in the infected group was downregulated (Extended Data Fig. 4f), the expression and location of non-muscle myosin IIA and phosphorylated myosin light chain (P-MLC) did not change, compared with non-infected cells (Fig. 2c-e, Extended Data Fig. 4e), indicating actomyosin-dependent contraction was required for VICM. Overall, VACV infection resulted in depolymerization of robust actin stress fibers but did not influence myosin-dependent contraction.

**Figure 2.**
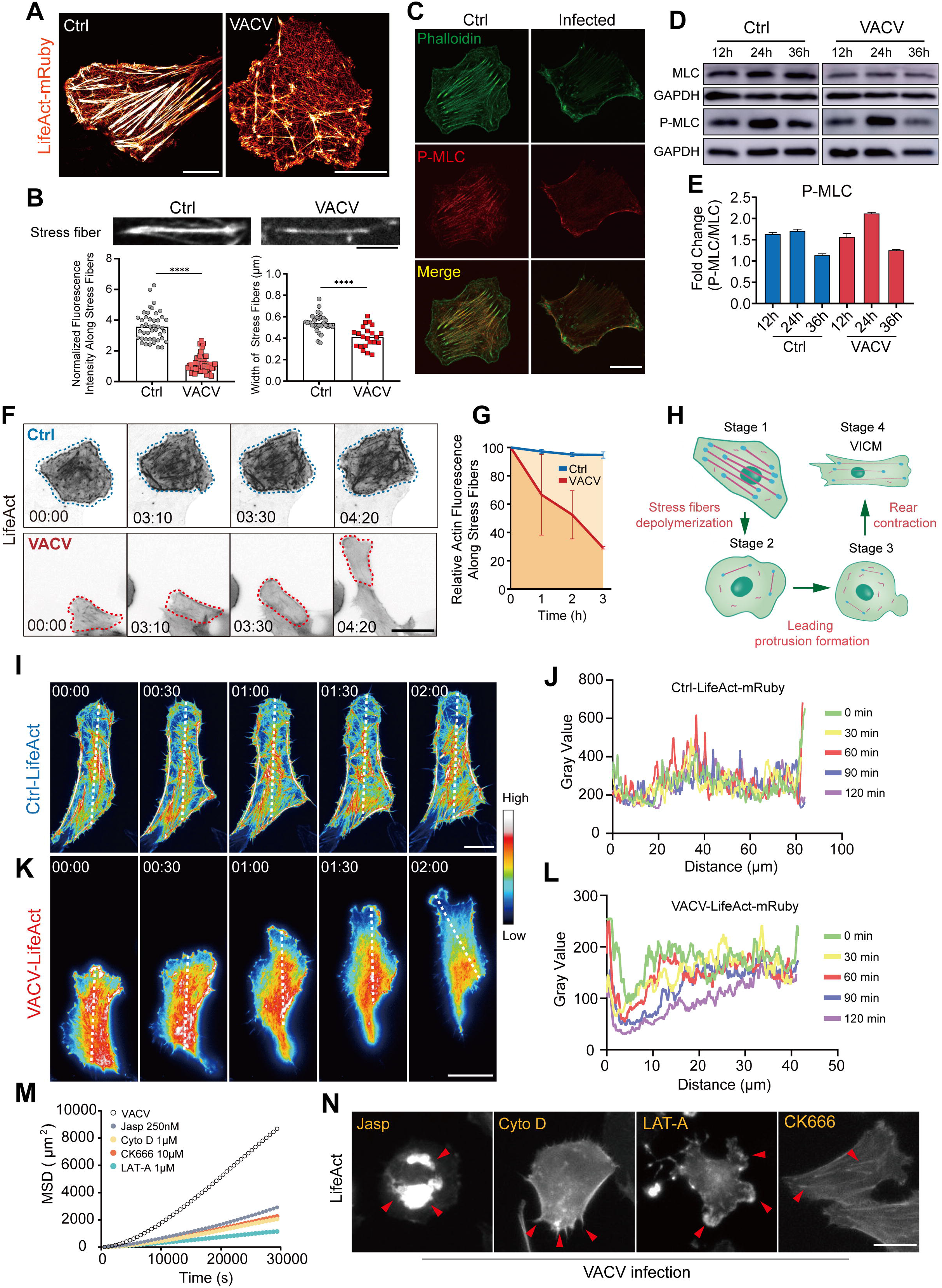
VICM is actin dependent. **a**, Representative super-resolution images of a control Vero cell (left) and a VACV-infected Vero cell (right) expressing LifeAct-mRuby. Scale bars represent 10 μm. **b**, Representative actin stress fiber images (top) and fluorescent quantitative (bottom left) and width (bottom right) of analysis stress fibers in control and VACV-infected cells by phalloidin staining. Scale bar represents 5 μm. **c**, Representative immunofluorescence images of F-actin (phalloidin, green) and p-MLC (red) in the control and VACV-infected cell. Scale bar represents 20 μm. **d**,**e**, Immunoblotting (d) and quantitative (e) analysis of MLC and p-MLC in Vero cells during VACV infection (n=3). **f**,**g**, Montage of frames from live-cell imaging (f) and relative fluorescent intensity analysis (g) of control and VACV-infected Vero cells expressing LifeAct-mRuby. The dotted lines represent the cell outlines (blue: control cells; red: VACV-infected cells). Scale bar represents 40 μm. **h**, Schematic diagram of the process of VICM accompanied by stress fibers depolymerization, leading protrusion formation, and rear contraction. **i**,**j**, Time-lapse actin heatmap (i) and plot profile of gray value (j) along white dotted line in a control cell in two hours. Scale bar represents 20 μm. **k, l**, Time-lapse actin heatmap (k) and plot profile of gray value (l) along white dotted line in a VACV-infected cell in two hours. Scale bar represents 20 μm. **m**,**n**, MSD (m) and representative images of actin cytoskeleton distribution (n) of Vero-LifeAct-mRuby cells during VACV infection after 250 nM jasplakinolide, 1 μM cytochalasin D, 10 μM CK666, and 1 μM latrunculin A treatment. Scale bar represents 20 μm. Data are mean ± SEM. *p<0.05, **p<0.01, ***p<0.001, ****p<0.0001 (unpaired t-test).

Live-cell imaging further confirmed robust actin stress fibers were gradually depolymerized during VICM compared to the control group (Fig. 2f, g and Supplementary Video 1). What’s more, we observed three main steps of VACV-induced migrating cells according to the dynamics of actin filaments and cell morphology (Fig. 2h, Extended Data Fig. 4g and Supplementary Video 2). Step 1: robust actin stress fibers depolymerization; Step 2: leading protrusion formation; Step 3: rear contraction and body translocation. High-resolution imaging showed actin signals were preferentially positioned at the front and back of migrating cells but had no tendency in non-infected cells during the observation time (Fig. 2i-l and Supplementary Video 3, 4). When VACV-infected cells were treated with actin assembly-related inhibitors (Jasplakinolide (Jasp), Cytochalasin D (Cyto D), CK666, and Latrunculin A (LAT-A)), VICM was significantly inhibited (Fig. 2m, Extended Data Fig. 5a). To confirm whether these inhibitors affected VACV infection, we tested virus infection by quantifying the EGFP fluorescence signal intensity. The statistics showed these inhibitors did not influence VACV infection except CK666 (Extended Data Fig. 5b, c). Meanwhile, VACV-infected cells exhibited different disrupted actin networks in the presence of these inhibitors, including Jasp-caused actin perinuclear aggregation, Cyto D-caused filopodia-like protrusions, and LAT-A-caused multi-protrusions (Fig. 2n and Supplementary Video 5). Collectively, VACV-induced actin dynamics together with myosin motor were crucial for VICM.

### RhoA signaling is dominant in regulating VICM

Time-resolved proteome analysis revealed that the Rho family of GTPases and its downstream effectors were downregulated during VACV infection (Fig. 3a, b). Similarly, the expression of associated genes was downregulated (Fig. 3c), indicating that VACV could manipulate the cytoskeleton by regulating these GTPases during VICM. Indeed, the expression of RhoA, Rac1, and Cdc42 were significantly downregulated when VICM occurred (Fig. 3d,e Extended Data Fig. 6a-c). In addition, RhoA is involved in protein-protein interaction (PPI) networks according to differentially expressed proteins (Fig. 3f). To test how RhoA, Rac1, and Cdc42 regulated VICM, we performed the inhibitor assays without influencing VACV infection (Extended Data Fig. 6d, e). The single-cell tracking-based analysis showed that Rho-associated protein kinase (ROCK) inhibitor Y27632 significantly suppressed VICM compared to Rac1 inhibitor NSC23766 (Fig. 3g, h, Extended Data Fig. 6f, g). However, Cdc42 inhibitor ML141 did not affect VICM. This suggested that RhoA played the most critical role in regulating VICM, followed by Rac1. Furthermore, the VACV-infected cells expressing dominant-negative RhoA-T19N and constitutively active RhoA-Q63L showed a lower migratory ability, indicating that VACV need proper RhoA activity to induce cell migration (Extended Data Fig. 6h). As a downstream effector kinase of RhoA, ROCK can regulate actin cytoskeleton and cell migration through phosphorylating MLC and focal adhesion kinase (FAK)^29,30^. We then examined the proteins related to the RhoA signal pathway after VACV infection. The gene and protein expressions of ROCK1 and FAK (phosphorylated or non-phosphorylated forms) were also downregulated during the time of VICM (Fig. 3a-e), although ROCK2 was not included (Fig. 3b-d). Moreover, FAK inhibitor Y15 gravely impaired VICM in terms of individual cell normalized trajectories, mean speed, and diffusion coefficient (Fig. 3i, j, Extended Data Fig. 6i). Altogether, VACV-manipulated RhoA signaling was responsible for VICM.

**Figure 3.**
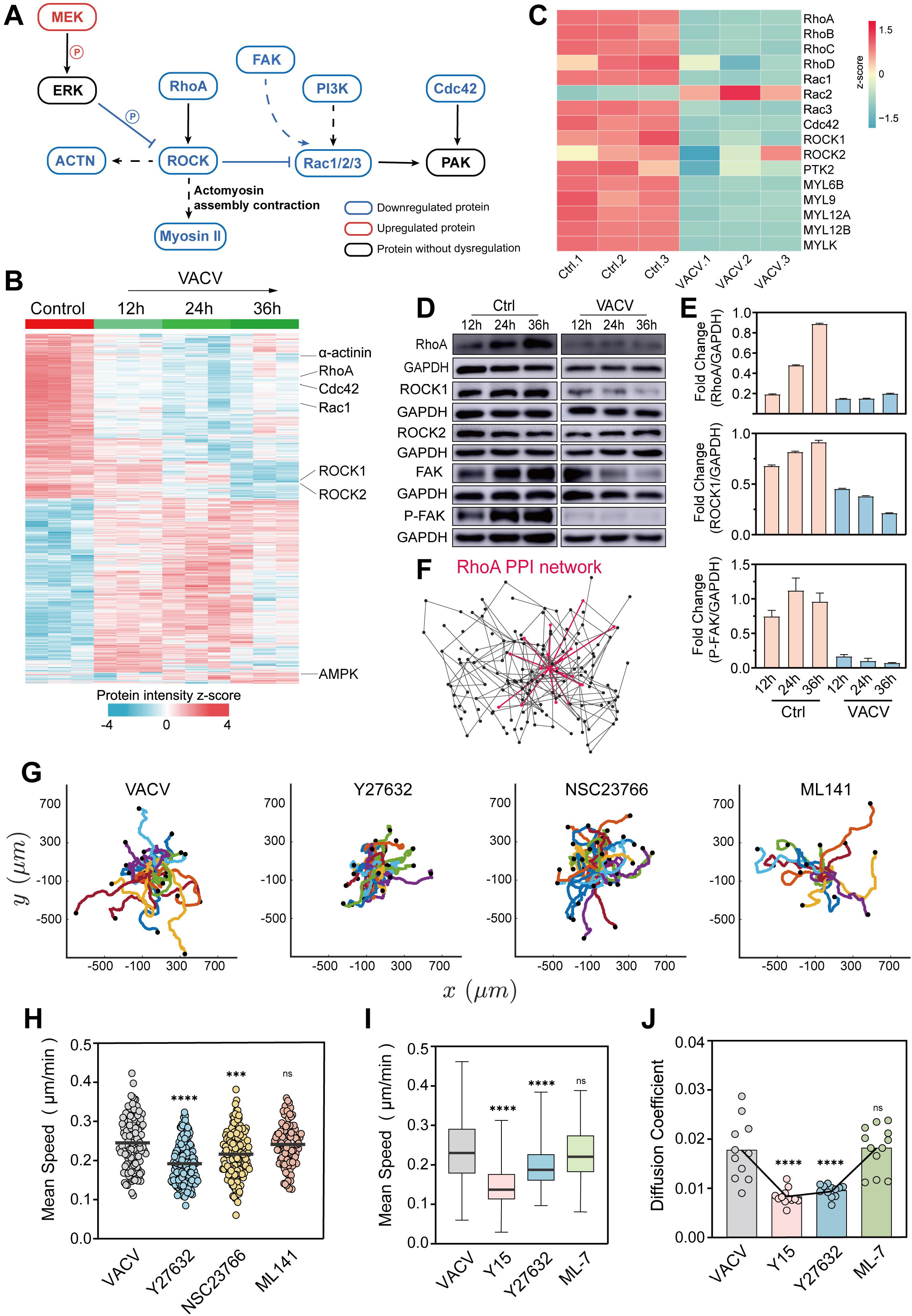
RhoA signaling regulates VICM. **a**, Proteomics-based RhoA signaling network enrichment during VACV infection. **b**, Heatmap of all differential proteins in Vero cells after VACV infection. Each column represents one biological replicate. **c**, RNA-seq-based gene expression level (FPKM) of small GTPase proteins and associated proteins during VACV infection. Each column represents one biological replicate. **d**,**e**, Immunoblotting (d) and quantitative analysis (e) of RhoA-related proteins in Vero cells during VACV infection (n=3). **f**, Protein-Protein Interaction (PPI) networks of DEPs. RhoA-associated PPI has been highlighted. **g**,**h**, Trajectory (g) and mean speed (h) of VACV-infected Vero cells after 80 μM Y27632, 250 μM NSC23766, and 500 nM ML141 treatment. **i**,**i**, Mean speed (i) and diffusion coefficient (j) of VACV-infected Vero cells after Y15 10 μM, Y27632 80 μM, and ML-7 10 μM treatment. Data are mean ± SEM. *p<0.05, **p<0.01, ***p<0.001, ****p<0.0001 (unpaired t-test).

### Inhibition of RhoA/ROCK promotes the depolymerization of robust actin stress fibers

Robust actin stress fibers in non-infected cells act as “the stabilizer” anchoring the cells in place. Before inducing cell migration, VACV needed to depolymerize the robust actin stress fibers in infected cells (Fig. 4a and Supplementary Video 6). We found only Y27632 treatment caused the depolymerization of actin stress fiber in non-infected cells, similar to the VACV infected cells (Fig. 4b, Extended Data Fig. 7a, b). This effect suggested that inhibition of RhoA/ROCK could disaggregate actin bundles. Western blot analysis showed that Y27632 can reduce the phosphorylation of MLC, FAK, and myosin phosphatase target subunit 1 (MYPT1) by directly inhibiting ROCK activity in non-infected cells (Extended Data Fig. 6c-h). By contrast, VACV infection also decreased the phosphorylation of FAK and MYPT1 by partially downregulating the expression of RhoA and ROCK1 (Extended Data Fig. 6c-h). In addition to viral F11L protein^13^, VACV can inhibit the RhoA/ROCK signal pathway in this manner to prepare for subsequent cell migration.

**Figure 4.**
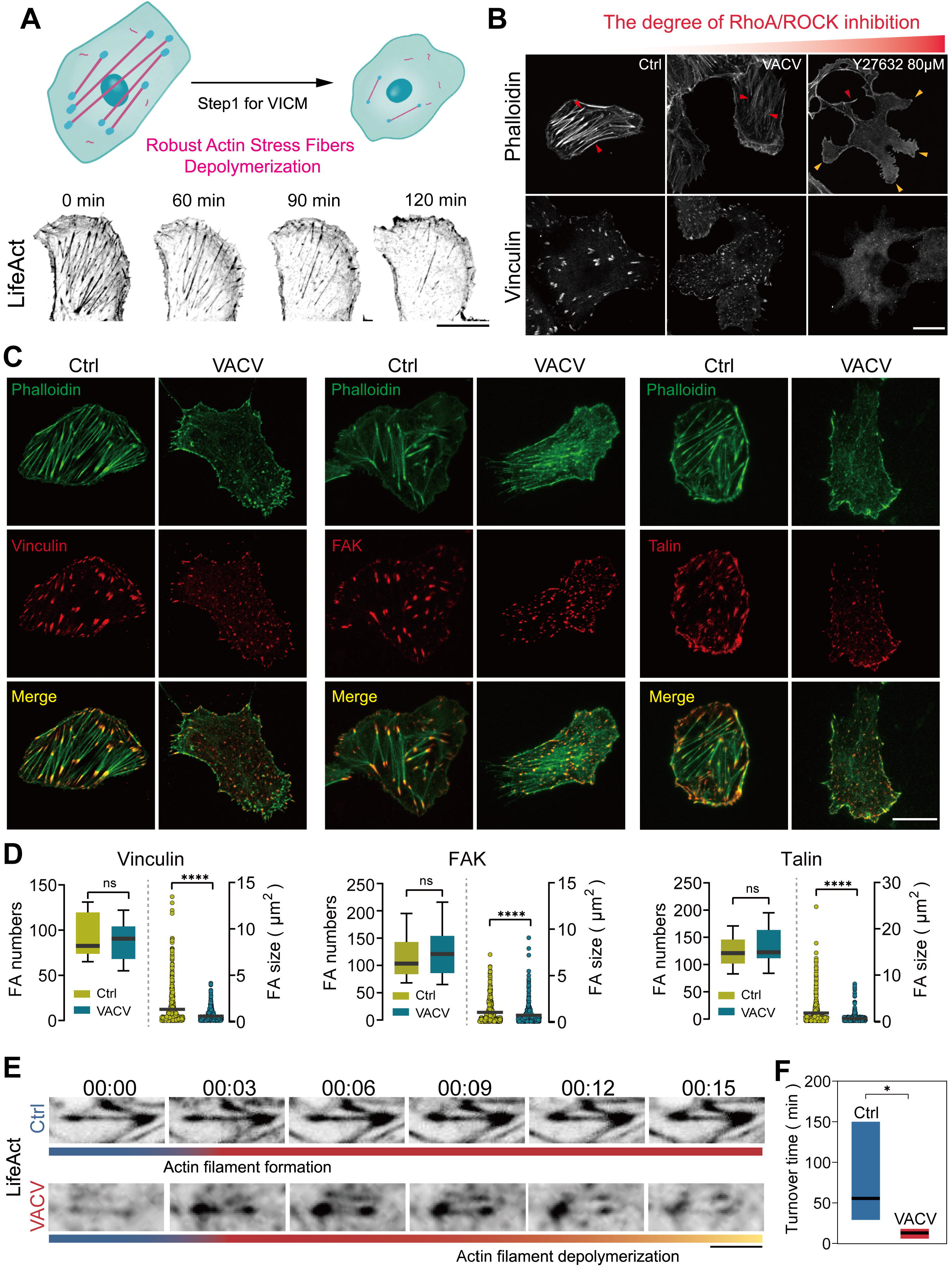
RhoA signaling regulates the stability of actin stress fiber during VICM. **a**, Schematic diagram of dynamics of actin stress fibers in a VACV-infected cell (top) and time-lapse images (bottom) of VACV-infected Vero cells expressing LifeAct-mRuby during actin stress fibers depolymerization. Scale bar represents 20 μm. **b**, Representative immunofluorescence images of F-actin (phalloidin; top) and Vinculin (bottom) in the control, Y27632-treated, and VACV-infected Vero cells. Red arrows indicate actin stress fibers and yellow arrows indicate the protrusions in the cells. Scale bar represents 20 μm. **c**, Representative immunofluorescence images of F-actin (phalloidin; green), vinculin (left, red), FAK (middle, red) and talin (right, red) in control and VACV-infected cells. Scale bar represents 20 μm. **d**, Quantitative analysis of FA numbers and size in control and VACV-infected cells according to immunofluorescence images of vinculin, FAK and talin. **e**,**f**, Time-lapse images (e) and turnover time (f) of actin stress fibers in control and VACV-infected cells. Scale bar represents 5 μm. Data are mean ± SEM. *p<0.05, **p<0.01, ***p<0.001, ****p<0.0001 (unpaired t-test).

High-resolution imaging showed almost all FAs with bigger plaque sizes were located at both ends of actin stress fibers in non-infected control cells (Fig. 4c). VACV-mediated RhoA/ROCK inhibition resulted in a smaller size of FAs and finer actin stress fibers compared to the control (Fig. 4b, c). Furthermore, nearly all mature FAs and stress fibers were lost in the cells at a higher degree of RhoA/ROCK inhibition by 80 μM Y27632 (Fig. 4b). Hence, we assumed VACV inhibited the maturation of FAs to regulate actin stress fibers dynamics. Similarly, Y15 (a potent FAK inhibitor) treatment not only inhibited the maturation of FAs but also led to depolymerization of actin stress fibers in non-infected cells, which further confirmed FAs dynamics influenced actin stress fibers assembly (Extended Data Fig. 7i). After VACV infection, FAs with small size exhibited immature characteristics accompanied by depolymerization of actin stress fibers (Fig. 4b-d). However, there was no difference in the number of FAs between the non-infected control cells and VACV-infected cells, as well as the expression of vinculin and talin (Fig. 4d, Extended Data Fig. 7j-l). These results suggested that VACV can also regulate the stability of actin stress fibers by affecting the maturation of FAs. What’s more, VACV-caused inhibition of RhoA/ROCK signaling accelerated the turnover of FA and actin stress fibers, which helped VACV coordinate the actin cytoskeleton during the time of VICM (Fig. 4e, f and Supplementary Video 7).

Taken together, VACV can depolymerize robust actin stress fibers and promote the turnover of finer actin stress fibers by inhibiting RhoA/ROCK/FAK signaling to prepare for VICM.

### VACV-regulated RhoA/ROCK promotes the leading protrusion formation

After the depolymerization of strong actin bundles, the leading protrusion with continuous extension was required for VACV-induced migrating cells (Fig. 5a and Supplementary Video 6). Live-cell imaging showed similar protrusion, but without continuous extension in the non-infected cells, around which a lot of actin stress fibers and FAs aggregated (Fig. 5b, Extended Data Fig. 8a and Supplementary Video 8). However, few actin stress fibers, and FAs gathered around the faster protrusions during VICM (Fig. 5c, Extended Data Fig. 8a,b and Supplementary Video 9). Moreover, FAs dynamic analysis showed FAs around VACV-induced protrusion were less stable (Fig. 5d and Supplementary Video 10, 11). Therefore, we supposed that the stress fibers, together with FAs, were important cellular components stabilizing extensive protrusion, and inhibition of RhoA/ROCK signaling promoted extensive protrusion generation. To test the hypothesis, we used 80 μM Y27632 treating Vero LifeAct-mRuby cells to inhibit the RhoA/ROCK signaling cascade. Intriguingly, non-infected cells not only displayed the depolymerization of actin stress fibers but also showed many dynamic protrusions after treatment, similar to the leading protrusion induced by VACV (Fig. 5e, top panel and Supplementary Video 12). This demonstrated that extensive protrusion was associated with inhibition of RhoA/ROCK. However, the number of protrusions in VACV-infected cells was generally less than four (Extended Data Fig. 8c, d). Because once the VACV-induced protrusion was formed, it would remain relatively stable unless the cell changed the direction of movement. In non-infected control cells, Y27632, not NSC23766 or ML141 treatment, can induce more than four short-time protrusions, which were random and unstable (Extended Data Fig. 8c, d). Based on these results, we considered that 80 μM Y27632 was too high, so that almost all stress fibers depolymerized. Serial dilution of Y27632 assay displayed 10 μM Y27632 could induce relatively stable protrusion formation in non-infected cells (Fig. 5e, middle panel and Supplementary Video 13). Additionally, inhibition of FAK by 10 μM Y15 can also induce dynamic protrusion in non-infected cells (Fig. 5e, bottom panel and Supplementary Video 14). Hence, as the result of moderate inhibition of RhoA/ROCK signaling, the extensive protrusion was stably generated (Fig. 5f). To further prove this hypothesis, we tested the ROCK activity in control, VACV-infected, and Y27632-treated cells. The activity in Y27632-treated cells was significantly inhibited, but there was no difference between non-infected control cells and VACV infected cells (Extended Data Fig. 8e). This may be because the difference in RhoA activity between non-infected and VACV-infected cells was too subtle to be detected by ROCK activity assay kit. Surprisingly, when we detected RhoA activity by the FRET sensor in the above three groups, we found activated RhoA signal was evenly distributed in non-infected control cells but located in protrusions in Y27632-treated cells (Fig. 5g). In other words, global inhibition of RhoA/ROCK activity can result in spontaneous activation of RhoA around the cell membrane where generated protrusion formation. In VACV-infected cells, high FRET efficiency appears at the leading protrusion, which suggested activated RhoA regulated VACV-induced protrusion formation. Moreover, super-resolved structured illumination microscopy (SIM) clearly observed the lamellipodia-like actin network in the protrusion. All protrusions were composed of crosslinking finer actin filaments demonstrating that the extensive protrusion formation required more fine actin networks rather than robust actin stress fibers (Extended Data Fig. 8f). Taken together, after overall inhibition of RhoA signaling, locally activated RhoA was responsible for the protrusion formation during VICM.

**Figure 5.**
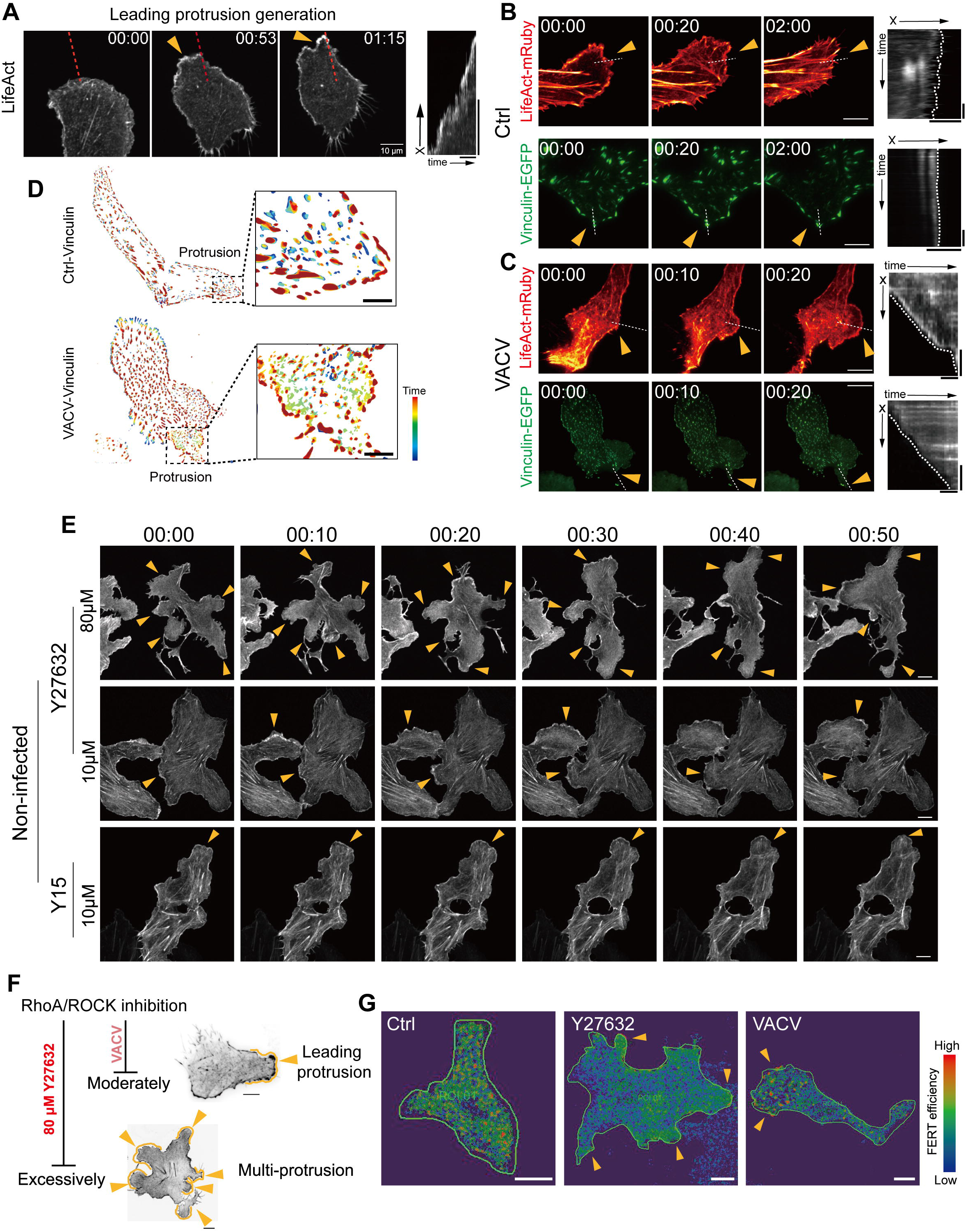
RhoA signaling regulates the protrusion formation during VICM. **a**, Time-lapse images (left) and kymograph (right) of VACV-infected Vero cells expressing LifeAct-mRuby during the protrusion formation. Horizontal bar (right) is 25 min and vertical bar is 10 μm. **b**,**c**, Time-lapse images (left) and kymograph (right) of Vero cells expressing LifeAct-mRuby (red) or vinculin-EGFP (green) with or without VACV infection around cell protrusion. Scale bars (left) represent 10 μm. Horizontal bars (right, top) are 5 μm and vertical bars (right, top) are 20 min. Horizontal bars (right, bottom) are 5 min and vertical bars (right, bottom) are 5 μm. **d**, Representative images of FA dynamics analyzed by Focal Adhesion Analysis Server (FAAS). The color represents the FA longevity (red: the longest lifetime; blue: the shortest lifetime). The black dotted line represents the VACV-induced protrusion in a cell. Scale bars represent 20 μm. **e**, Time-lapse images of Vero cells expressing LifeAct-mRuby after 10 μM or 80 μM Y27632 and 10 μM Y15 treatment. Yellow arrow indicates the cell protrusion. Scale bars represent 10 μm. **f**, Schematic diagram of the relationship between RhoA/ROCK inhibition (VACV or Y27632 80 μM) and the protrusion in the cells. Yellow arrows indicate the protrusions in the cells. Scale bars represent 10 μm. **g**, Representative FRET images of Vero cells expressing RhoA2.G (RhoA sensor) with Y27632 or VACV treatment. The color represents the FRET efficiency. (blue: the lowest efficiency; red: the highest efficiency). Scale bars represent 10 μm.

### VACV-modulated RhoA/ROCK activity regulates cell contraction

Coordinating with the leading edge protrusion, myosin-mediated rear contraction in migrating cells contributed to cell translocation (Fig. 2h). The molecular switch that regulates actomyosin contraction is the MLC phosphorylation controlled by myosin light chain kinase (MLCK) and ROCK^31,32^. MLC maintained high phosphorylation levels during the time of VICM (Fig. 2d, e), which indicated that myosin-mediated contractile force was indispensable. On the one hand, we found the gene and protein expression of MLCK in VACV-infected cells was downregulated (Fig. 3c and 6a), suggesting that MLCK-dependent MLC phosphorylation was disrupted by VACV. When we inhibited MLCK with ML-7 or ML-9, we found these inhibitors had no effect on VICM (Fig. 3i, j, Extended Data Fig. 6i and 8g-j). These results implicated that VACV eliminated the effect of MLCK by downregulating its expression. Furthermore, phosphorylation of MLC was not influenced by ML-7 in VACV-infected cells (Extended Data Fig. 8k, l), but was inhibited in non-infected cells, which confirmed that VACV infection inhibited the function of MLCK. On the other hand, inhibition of ROCK can remarkably downregulate the phosphorylation of MLC in VACV-infected cells (Fig. 6b), that is, RhoA/ROCK regulated phosphorylation of MLC assuring rear contraction of VACV-induced migrating cells. Meanwhile, time-lapse images and kymographs showed that VACV-induced cell contraction was consistent with the direction of cell migration (Fig. 6c, d and Supplementary Video 15). In contrast, spontaneous contraction of the cell periphery was bidirectional in non-infected cells. Further inhibiting ROCK by Y27632 in the virus-infected cell can destroy VACV-induced contraction because the originally contracted cell part became a protrusion and extended back (Fig. 6e and Supplementary Video 16). Therefore, VACV-regulated RhoA/ROCK signaling was responsible for the rear contraction of moving cells through phosphorylating MLC.

**Figure 6.**
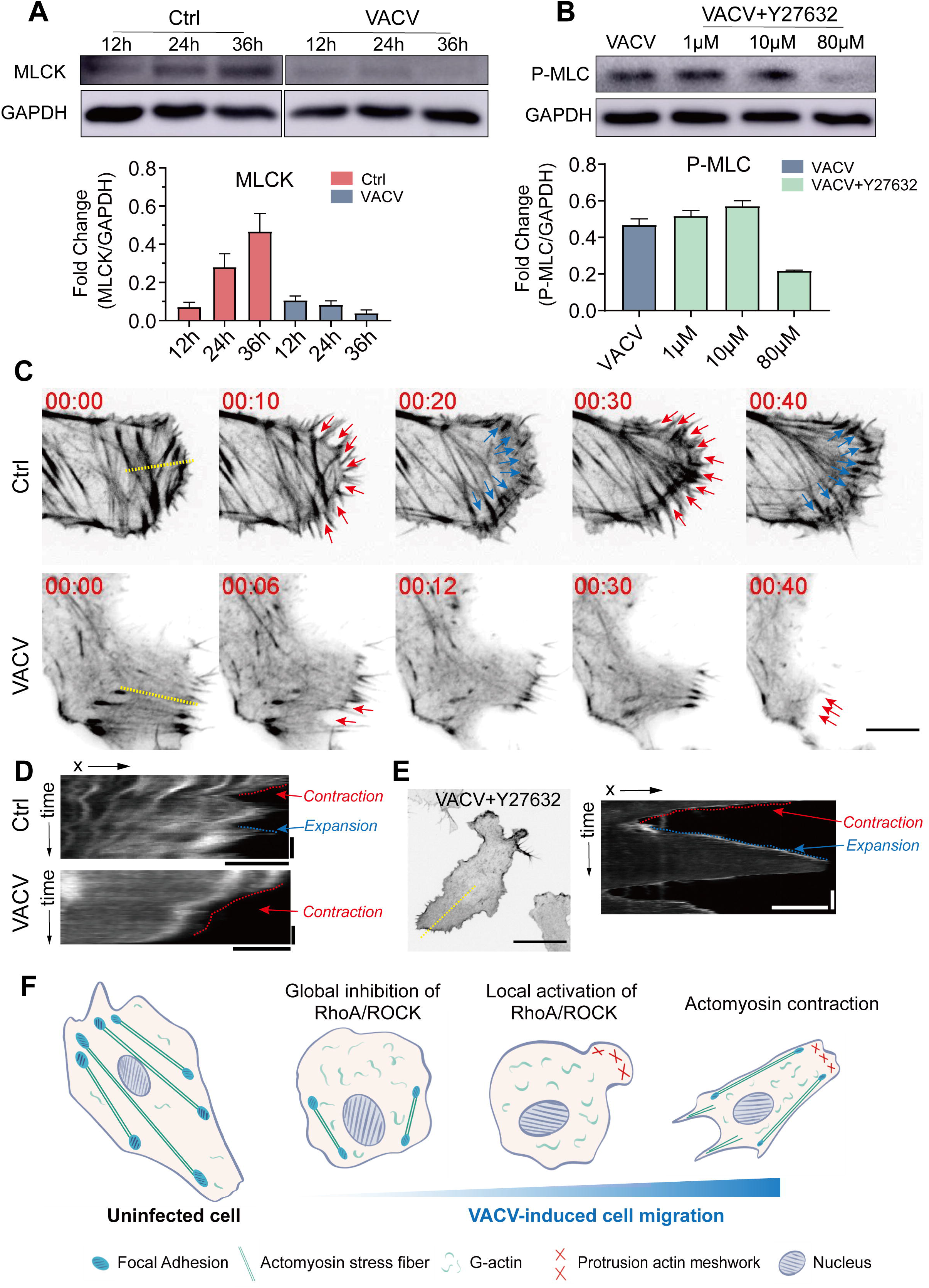
RhoA signaling regulates the cell contractility during VICM. **a**, Immunoblotting (top) and quantitative (bottom) analysis of MLCK in Vero cells with or without VACV infection (n=3). **b**, Immunoblotting (top) and quantitative (bottom) analysis of p-MLC in VACV-infected Vero cells with different Y27632 concentrations (n=3). **c, d**, Time-lapse images (c) and kymograph (d) of contractile part in Vero cells expressing LifeAct-mRuby with (bottom) or without (top) VACV treatment. Highlighted in red indicates cell part contraction and highlighted in blue means cell part expansion. Scale bar (c) represents 10 μm. Horizontal bars (d) are 5 μm and vertical bars (d) are 10 min. **e**, Representative images (left) and kymograph (right) of Y27632-treated Vero-LifeAct-mRuby cells with VACV infection. Scale bar (left) represents 20 μm. Horizontal bar (right) is 10 μm and vertical bars (right) is 30 min. f, A model for VACV-induced cell migration.

In conclusion, VACV infection can precisely regulate RhoA/ROCK signaling to depolymerize robust actin stress fibers, generate leading protrusion, and maintain actomyosin-mediated contraction (Fig. 6f). Through this process, VACV is able to manipulate cell migration spreading the nascent virus.

## Discussion

Virus infection can alter the morphology and phenotype of cells, which is response for viral transmission, pathogenesis, and carcinogenesis^14,33^. For example, the epithelium infected by severe acute respiratory syndrome coronavirus 2 (SARS-CoV-2) became a mesenchymal state via up-regulation of ZEB1 and AXL^34^. In our study, we found the EMT-like process in VACV-infected cells due to losing epithelial cell-cell adhesions and acquiring migratory properties (Fig. 1a-f). The downregulation of E-cadherin, a hallmark of EMT, is generally associated with activating canonical Wnt/β-catenin signaling, which regulates cadherin-related transcription factors and cadherin-mediated cell adhesion^35,36^. However, the inhibition of Wnt/β-catenin signaling did not affect VACV-induced cell migration after the EMT-like process (unpublished data), indicating that VACV-induced EMT was Wnt-independent. Transcription factors play an essential role in inducing EMT^7^. We found that transcription factors of the SNAIL, ZEB, and TWIST families were upregulated in VACV-infected cells (Fig. 1d), further increasing a VACV-induced EMT score. Given that Smad4-dependent induction of PAI-1 and Smad4-independent induction of Snail and Snail2 were all involved in VACV-induced EMT^20^, we propose that VACV-induced EMT-like cell transformation requires multi-pathways-mediated transcription factors activation. Further investigation can elaborate on the roles of different transcription factors and determine which pathway predominates in VACV-induced EMT.

VACV-infected cells gain migratory capacities during the EMT-like biological process. Since numerous nascent viruses are wrapped inside the cell, VICM is a good choice for the virus to avoid immune system defense such as antibody neutralization. In addition, VACV infection can impair the migratory capacity of skin dendritic cells from the infection sites to draining lymph node^37^, which is also a potential way of immune escape for the vaccinia. Besides cell-free transmission, viruses can also spread the emerging virus through cell-contacts-based cell-to-cell transmission, subdivided into the virological synapse-based and trans-infection-based manner^38,39^. For example, virus particles in HIV-infected CD4^+^ T cells can be released to mucosal epithelium through virological synapses and then transferred into a microenvironment where macrophages are susceptible to infection^40^. Cell-to-cell transmission, including VICM, is a more efficient than cell-free transmission to spread the virus and overcome physiologic barriers^41–43^. Thus, viruses use different strategies for their survival and distribution in the host. A comprehensive understanding of cell-to-cell virus transmission can be conducive to figuring out the exact pathogenesis of virus infection and provides novel targets for intervention, especially for SARS-CoV-2.

VACV-regulated cell migration requires the coordination of cytoskeletal reorganizations and signal pathways in the host. We found that the actin cytoskeletal rearrangement played a dominant role in VICM. Before VACV infection, strong actin stress fibers with mature FAs formed stable cytoskeletal structures and maintained cell shape. VACV first affected the stability of strong actin stress fibers and FAs by inhibiting RhoA/ROCK signaling, preparing to induce cell migration. Significantly, we found that VACV infection did not depolymerize all stress fibers but retained some finer actin bundles with a short turnover time. Then the more dynamic, delicate actin stress fibers in VACV-infected cells coordinated the cell migration. We speculate that down-regulated expression of RhoA is another way of inhibiting RhoA/ROCK signaling except for viral F11L protein^13,25^. VACV-mediated RhoA/ROCK inhibition is precisely regulated because a high degree of RhoA/ROCK inhibition results in loss of all stress fibers, which is not conducive to VICM.

Surprisingly, RhoA signaling plays a dominant role in VICM. In the canonical description of cell motility, RhoA is involved in tail retraction, FA dynamics, and cell protrusion^44,45^, and Rac1 is required for leading-edge protrusion and lamellipodia formation^46^. In mutually antagonistic relationships, RhoA and Rac1 play a dominant role in different cell migration modes under different extracellular cues^44,47,48^. We found the expression of RhoA, Rac1, and Cdc42 in VACV-infected cells was dramatically downregulated, and the complex interplay of these small GTPases may be broken. Consequently, not suppressed by Rac1 activity, RhoA activity was mainly engaged in VICM. RhoA/ROCK signal pathway was entirely responsible for the depolymerization of robust actin stress fibers, the protrusion at the leading edge, and rear contraction, indicating two different ROCK isoforms (ROCK1 and ROCK2) might play different roles in regulating VICM. Further research can focus on the individual effect of ROCK1 or ROCK2 during VICM. In addition, we found that Y27632 treatment resulted in extensive protrusions in non-infected cells, similar to VACV-induced leading-edge protrusions. As RhoA-GTP accumulated near the protrusions (Fig. 5g), we speculate a spontaneous, membrane-bound RhoA activation is responsible for protrusion formation after global ROCK inhibition. The molecular mechanism behind it remains to be discovered. Overall, our findings mainly focused on the dynamics of cellular components during VICM and did not disclose which viral elements participated in VICM. Elaborating on both cellular and viral components involved in VICM can determine the mechanism of VICM and provide a potential possibility for manipulating actin cytoskeleton and even cell migration.

## Methods

### Reagents and plasmid

Deionized (DI) water (18.2 MΩ·cm) was produced from a Milli-Q system (Millipore, Bedford, MA, USA) and used in all experiments. HPLC grade acetonitrile (ACN) and methanol, as well as analytical grade acetone and hydrochloric acid (HCl, 37%) were purchased from Sinopharm Chemical Reagent Co., Ltd. (Shanghai, China). Analytical grade formic acid was purchased from J&K Scientific Ltd. (Beijing, China). Iodoacetamide (IAA), trizma base, urea, sodium dodecyl sulfate (SDS), ammonium bicarbonate (NH_4_HCO_3_) and caffeine were purchased from Sigma-Aldrich (St. Louis, MI, USA). Bond-breaker TCEP solution (0.5 M) and protease inhibitor cocktail (100×, EDTA-free) were purchased from Thermo Fisher Scientific (Rockford, USA). Trypsin used for protein digestion was bought from Hualishi Technology Co., Ltd. (Beijing, China). Trypsin-EDTA solution (0.25%, with phenol red) used for cell digestion and phosphate buffered saline (PBS, 1×) were purchased from Solarbio Science & Technology Co., Ltd. (Beijing, China). pLL(20)-g[3.5]-PEG(2) (pLL-PEG) was purchased from SuSoS. Alexa Fluor™ 594 or 647 Phalloidin was purchased from Invitrogen. CK666 was purchased from Tocris. Y27632 was purchased from Millipore. Jasplakinolide, cytochalasin D, latrunculin A, Y15, tetracycline, ML-7 hydrochloride, and ML-9 were purchased from MedChemExpress. NSC 23766 trihydrochloride and ML141 were purchased from TergetMol. Fibronectin, Hoechst 33342, monoclonal anti-α-Tubulin antibody, and anti-vinculin antibody were purchased from Sigma-Aldrich. MYLK polyclonal antibody and myosin light chain polyclonal antibody were purchased from Proteintech. ROCK1 or ROCK2 rabbit mAb, GAPDH rabbit mAb, FAK rabbit mAb, phospho-FAK rabbit mAb, Rac1/2/3 rabbit mAb, Cdc42 rabbit mAb, E-Cadherin rabbit mAb, N-Cadherin rabbit mAb, phospho-myosin light chain 2 rabbit mAb, Vimentin rabbit mAb, ZO-1 rabbit mAb, phospho-MYPT1 antibody, and Myosin IIa antibody were purchased from CST. Anti-RhoA antibody, anti-Talin 1 and 2 antibody, anti-non-muscle Myosin IIB antibody, anti-cytokeratin 14 antibody, and anti-Vaccinia Virus antibody were purchased from Abcam. Anti-MYPT1 Antibody was from Santa Cruz. The plasmids (pLenti Lifeact-mRuby2 BlastR, pLentiRhoA2G, tetO-FUW-EGFP-RhoA-Q63L, tetO-FUW-EGFP-RhoA-T19N) were purchased from Addgene. psPAX2 and pMD2.G were generously given by Fa-Xing Yu (Fudan University). pLenti vinculin-EGFP was generously given by Congying Wu (Peking University).

### Cell culture and sample preparation

Human embryonic kidney 293T cells, kindly provided by Guo-Liang Xu (Fudan University), and Vero cells were cultured in DMEM (GIBCO) supplemented with 10% fetal bovine serum (GIBCO), 100 U/mL penicillin and streptomycin (GIBCO) at 37 °C and 5% CO_2_. Vero cells expressing LifeAct-mRuby or vinculin-EGFP were cultured in DMEM supplemented with 10% fetal bovine serum, 100 U/mL penicillin and streptomycin, 500 ng/mL blasticidin S (Millipore) or puromycin (Millipore) at 37 °C and 5% CO_2_. All samples were incubated with 200 ng/mL Hoechst 33342 for 1h at 37 °C before live-cell imaging.

### Viruses

For virus stock, Vero cells monolayers were infected with VACV for 72h in DMEM containing 2% FBS. After three freeze-thaw cycles and centrifugation at 12,000 rpm 4 °C for 15 min, the lysates were collected and stored at −80 °C. The VACV titers were quantified according to The TCID50 (Median Tissue Culture Infectious Dose) assay.

### Analysis of cell trajectories

For cell motility analysis, the nucleus images labeled by Hoechst were tracked using TrackMate plugin in FIJI. Briefly, the calibrated-image sequences were detected by Log detector and were tracked by simple LAP tracker. The spots in tracks statistics were further analyzed by self-wrote code in Matlab to obtain the normalized cell trajectories, mean speed, MSD, and diffusion coefficient. The velocity of migratory cells was calculated according to the displacement of the migratory cells between image sequences divided by timelapse. The mean square displacement (MSD) was calculated as the following Equation (1).

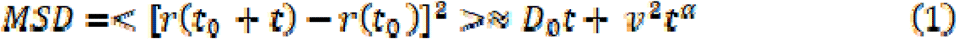

where υ is the mean speed of migratory cell, *D*_0_ is the thermal diffusion coefficient of the cell, is the time dependent position of migratory cell, r(*t)* and *t* is the timelapse of the position.

### Transwell-based virus spread assay

Vero cells at a density of 2×10^5^ cells/mL were seeded on 24-wells plates. When the cells were confluent in a monolayer, the anti-vaccinia virus antibody at a final concentration of 80 μg/mL or PBS was added to the culture medium. Transwells with 8 μm polyester membrane(BD Falcon)were incubated in DEME at 37 °C for 1h and placed on 24-wells plates. Viral supernatant with or without VACV-infected Vero cells was added to the upper compartment, and the EGFP fluorescence of the cells in the lower compartment was detected at 48 or 72 h.p.i.

### RNA-seq analysis

Vero cells were seeded at a density of 2×10^6^ cells/mL on 60 mm dishes. The non-infected or VACV-infected cells (at MOI 1) were treated with 1 mL TRIzol to extract RNA after washing three times with PBS. RNA library sequencing was performed on the Illumina HiseqTM 2500/4000 by Gene Denovo Biotechnology Co., Ltd (Guangzhou, China). Bioinformatic analysis was performed using Omicsmart, a real-time interactive online platform for data analysis (http://www.omicsmart.com).

### Proteomic analysis

Vero cells infected with VACV after 12 h, 24 h, and 36 h were harvested by trypsin-EDTA free digestion as well as the cells without treatment. Then the cells were washed with 1× PBS and centrifuged at 1000 rpm, 4 °C three times to gain cell pellets (about 6×10^5^ cells per plate) after the removal of supernatants. Cell pellets were rapidly transferred into a 1.2 mL Eppendorf tube. Subsequently, 400 μL of lysis buffer (20 mM Tris, 8 M Urea, 1% SDS, HCl, pH = 8∼9) and 4 μL of 100× protease inhibitor cocktail was added for cell lysis. After a thorough cell disruption by ultrasonication for 10 min, the supernatant was obtained by centrifuging at 15,000 rpm, 4 °C and stored at −80 °C. Pierce BCA protein assay (Thermo Fisher Scientific, Rockford, USA) was utilized to quantify protein concentrations. After protein quantification, 150 μg of protein was carbamidomethylated using 10 mM TECP for 1 h under 37 °C and 40 mM IAA (final concentration) for 45 min in the dark at 25 °C in 25 mM ammonium bicarbonate (ABC) solution with the protein concentration of 1∼2 μg/μL. Six times volumes of acetone (pre-cooled under 20 °C) was added to precipitate protein pellets. Furthermore, proteins pellets were centrifuged, washed with 90% acetone three times, and digested by adding 4 μL of trypsin in a ratio of trypsin: protein=1:50 (w/w) and incubated at the speed of 600 rpm/h overnight at 37 °C.

After digestion overnight, the supernatant was desalted with desalting columns (MonoSpin C18, GL Science, Inc, Tokyo, Japan) using the standard protocol in the kit. The obtained supernatant was dried in a vacuum drying system (LNG T98 vacuum dryer, Taicang Huamei Biochemical Instrument Factory, Jiangsu, China) at room temperature for 1 h. Dried peptides were resuspended in DI water and then quantified using Pierce Quantitative Colorimetric Peptide Assay (Thermo Fisher Scientific, Rockford, USA).

### Live-cell imaging

Time-lapse images were acquired with 20×, 40×, or 100× objectives using either a DMi8 inverted microscope (Leica) equipped with a ORCA-Flash4.0 V3 Digital CMOS camera (Hamamatsu) controlled by MetaMorph software (Universal Imaging) or a spinning disk confocal microscopy (Yokogawa, CSU-W1, Nikon) equipped with a prime 95B camera (Teledyne Photometrics) controlled by NIS-Elements software (Nikon) in a cage incubator (Okolab). Image analysis was performed using FIJI software.

### Multi-SIM super-resolution microscopy

TIRF-SIM images of Vero-LifeAct-mRuby cells were acquired on the Multimodality Structured Illumination Microscopy (Multi-SIM) system equipped with solid-state single mode lasers (488 nm, 561 nm, 640 nm) and a sCOMS (Complementary Metal-Oxide-Semiconductor) camera (ORCA-Fusion C15440-20UP, HAMAMATSU). To obtain optimal images, immersion oils with refractive indices of 1.518 were used for Vero-LifeAct-mRuby cells on glass coverslips. Time-lapse images were acquired with the 100X 1.49NA Objective (Nikon) from the specimens incubated in a stage top incubator (Okolab). The microscope is routinely calibrated with 100□nm fluorescent spheres to calculate both the lateral and axial limits of image resolution. SIM images were reconstructed with the following settings: pixel size 30.6□nm; channel-specific optical transfer functions; Wiener filter constant 0.01 for TIRF-SIM mode. Then the reconstructed SIM image was denoised with total variation (TV) constraint. Pixel registration was corrected to be less than 1 pixel for all channels using 100□nm fluorescence beads.

## Acknowledgments

This study was supported by grants from the National Natural Science Foundation of China (Nos. 31870978, 21934001 and 22274026). We thank Matthieu Piel (Institut Curie) for comments on the manuscript, Fa-Xing Yu (Fudan University) and Congying Wu (Peking University) for sharing plasmids, Qun Li (Fudan University) for helping in RNA-seq analysis.

## Author contributions

W.L., J.-Y.L., Y.-J.W. : Conceptualization, Investigation, Data analysis, Writing – original draft. X.-X.X, Y.-C.C., S.-X.Y., X.-W.X., X.-Z.C., M.S., X.H., D.L., T.Y. : Methodology, Data analysis. Y.J., H.G., Z.-J.C., P.M., X.H., G.-L.X., D.-W.P. : Writing – review & editing. Z.-L.Z., B.L. : Supervision, Writing – review & editing, Funding acquisition. Y.-J.L. : Conceptualization, Supervision, Writing – review & editing, Funding acquisition. All co-authors have seen and approved the manuscript.

