## Supplementary material for "Vaccinia virus induces EMT-like transformation and RhoA-mediated mesenchymal migration": Supplemetal materials

**Methods**

**Lentivirus production and Lentivector transductions**

Lentiviral particles were produced from 293T cells. Briefly, 293T cells were seeded in a 6-well plate. When cell confluency reached 50-60 %, the cells were transfected with 1.5 μg of psPAX2, 0.5 μg of pMD2.G, and 2 μg of a lentiviral vector plasmid (pLenti Lifeact-mRuby2 BlastR, pLentiRhoA2G, tetO-FUW-EGFP-RhoA-Q63L, tetO-FUW-EGFP-RhoA-T19N, or pLenti Vinculin-EGFP) using LipoFiter 3.0 (Hanbio, Shanghai, China) per well. The medium was changed to warm DMEM with 10 % FBS at 24 h post-transfection, and the supernatant was harvested twice at 48 and 72 h post-transfection.

For lentivector transductions, Vero cells at a density of 1×10^4^ cells/mL on 6 well cell culture plates were infected with 1 mL viral supernatant and were selected by 2 μg/mL puromycin, tetracycline or blasticidin S at 72 h post-infection for several days.

**Western blot**

The cells with different treatments were removed from culture plates by cell scrapers (NEST) and resuspended in RIPA buffer (ThermoFisher) containing a protease inhibitor cocktail (ThermoFisher). After centrifugation at 12,000 rpm 4 °C for 10 min, the supernatants were diluted in NuPAGE™ LDS（Invitrogen）and boiled for 15 min. Protein samples were isolated on 8%-15% SDS–PAGE acrylamide gels and transferred onto PVDF membranes by wet electrophoretic transfer. Membranes were blocked with 5% BSA (Sigma) or nonfat milk (CST) in PBST containing 0.1% Tween-20 (Sigma) for 1 h at room temperature, followed by primary antibodies incubation at 4 °C overnight and PBST washing. Then, the membranes were incubated with secondary antibodies in PBST at room temperature for 1h and washed with PBST. ECL signals were detected by GE Amersham Imager 600.

**Immunofluorescence**

The cells with different treatments were seeded on 35 mm petri dishes coated with 25 μg/mL FN, fixed with 4% paraformaldehyde at room temperature for 15 min, and permeabilized in 0.25% Triton X-100 in PBS for 8 min. After washing in PBS, the cells were incubated with 3% BSA in PBS for 1h at room temperature, followed by primary antibodies incubation at 4 °C overnight or Alexa Fluor 488, 594, or 547-conjugated secondary antibodies for 1 h at room temperature with extensive PBS washing. Then cells were labeled with Alexa Fluor 488, 594, or 547-conjugated phalloidin and placed on a Nikon microscope for image capture.

**Drug treatment**

Vero cells expressing LifeAct-mRuby or vinculin-EGFP at a density of 1×10^5^ cells/mL were seeded on glass-bottom 6-well plates or 35mm petri dishes coated with 25 μg/mL fibronectin and infected with VACV at MOI 1. The drugs at a final concentration of 250 nM jasplakinolide, 1 μM cytochalasin D, 10 μM CK666, 1 μM latrunculin A, 80 μM Y27632, 250 μM NSC23766, 500 nM ML141, 10 μM Y15, 10 μM ML-7, and 20 μM ML-9 were added into VACV-infected cells at VACV 8 h.p.i. and were maintained all the time during image acquisition.

**Kymograph**

All kymographs were produced with FIJI software. Briefly, a line in the region of interest (ROI) was drawn, and the kymograph was generated by the Multi Kymograph tool.

**ROCK activity kit**

ELISA-based 96-well ROCK activity assay kit (Cell Biolabs) was used to measure the activated ROCK activity in non-infected, Y27632-treated, and VACV-infected cells. Briefly, the cells were lysed in lysis buffer containing 50 mM Tris-HCl, pH 7.5, 150 mM NaCl, 1 mM 2-glycerophosphate, 1% TritonX-100, 1 mM EDTA, 1 mM EGTA, 1 mM Na_3_VO_4_ and proteinase inhibitors and the lysates were centrifuged at 12,000 rpm 4 °C for 10 min and the supernatants were collected. Then, cell lysate samples or active ROCK-II positive control were added to the MYPT1-coated wells to incubate at 30 °C for 30-60 minutes with gentle agitation, followed by anti-phospho-MYPT1 (Thr696) antibody and HRP-conjugated secondary antibody incubation with extensive washing. Finally, substrate solution was added to each well, including the blank wells to incubate at room temperature for 5-20 minutes with gentle agitation, and then stop solution was added to each well. Read the absorbance of each microwell on a spectrophotometer using 450 nm as the primary wavelength.

**FRET assay**

Vero cells expressing RhoA2G sensor at a density of 1×10^5^ cells/mL were seeded on 60 mm dishes and treated with VACV or 80 μM Y27632. The FREF signal of the RhoA2G sensor was detected by using Leica TCS SP8 LAS AF. The imaging condition (CFP excitation 458 nm; emission 462-510 nm; YFP excitation 514 nm; emission 518-580 nm) was set up and the single-cell outlines as ROI were drawn for acceptor photobleaching. The images of the donor and acceptor were acquired for FRET efficiency calculation.

**Micropatterning assay**

The clean glass coverslips were activated in a plasma chamber (Harrick Plasma, Ithaca, NY, USA) for 1 min, modified with 0.1 mg/mL pLL-PEG in 10 mM pH 7.4 HEPES buffer at 4 °C overnight, washed in PBS three times, and then rinsed in H_2_O three times. Crossbow-shaped mask was cleaned with isopropanol, well-dried, and treated in UVO-Cleaner (Jelight Company) for 5 min. Then 3 μL H_2_O was dropped on the metal side of the mask where the coverslip with 0.1 mg/mL pLL-PEG was placed on. The mask with the coverslip was treated with a deep UV lamp for 1 min. After washing in H_2_O three times, the coverslip was modified with 25 μg/mL of fibronectin or Alexa Fluor 488-labeled fibronectin at room temperature for 1h. Vero cells at a density of 2×10^5^ cells/mL were inoculated on the micropatterning coverslips, incubated at 37 °C for 30 min, and washed with DMEM three times to remove the nonadherent cells.

**Focal adhesion dynamics analysis**

Vero cells expressing Vinculin-EGFP were seeded at a density of 5×10^4^ cells/mL on 35 mm petri dishes coated with 25 μg/mL FN. After VACV infection at MOI 1, the dishes were transferred in a cage incubator (Okolab) on a total internal reflection fluorescence microscopy (TIRF, Nikon) equipped with a prime 95B camera (Teledyne Photometrics). The timelapse images were acquired with NIS-Elements software (Nikon) and were processed for FAs dynamics by the Focal Adhesion Analysis Server (FAAS) (<http://faas.bme.unc.edu>).

**Nano-UPLC-MS/MS analysis**

The iRT kit (Ki3002, Biognosys AG, Switzerland) was added to all of the samples to calibrate the retention time of extracted peptide peaks. Then, all samples were analyzed by online nano flow liquid chromatography tandem mass spectrometry performed on an EASY-nanoLC 1200 system (Thermo Fisher Scientific, MA, USA) connected to a Orbitrap Q-Exactive HF-X mass spectrometer (Thermo Fisher Scientific, MA, USA). 3 μg peptides of each sample were loaded and analyzed in data independent acquisition (DIA) mode. The survey of full scan MS spectra (m/z 350-1500) was acquired in the Orbitrap with 60,000 resolution. All precursor ions were entered into collision cell for fragmentation by higher-energy collision dissociation (HCD), the collision energy was 28. The MS/MS resolution was set at 30,000.

**DIA Raw Data analysis**

Raw Data of DIA were processed and analyzed by Spectronaut 15 (Biognosys AG, Switzerland) with default settings and retention time prediction type was set to dynamic iRT. Searched database was downloaded from the UniProt database (https://www.uniprot.org/taxonomy/60711, June 28, 2021, containing 19,229 proteins). Q-value (FDR) cutoff on precursor level was 1% and protein level was 1%. All selected precursors passing the filters are used for quantification. MS2 interference will remove all interfering fragment ions except for the 3 least interfering ones. The average top 3 filtered peptides which passed the 1% Q-value cutoff were used to calculate the major group quantities.

**PRM analysis**

The control group (without treatment) and one experimental group (with 24 h infection) were selected to launch the parallel reaction monitoring (PRM) analysis. The peptides of two groups, including three biological replicates, were taken and mixed equally to make a mixture sample, then re-dissolved in solvent A (0.1% formic acid in water) and analyzed by on-line nanospray LC-MS/MS on Orbitrap Fusion Lumos mass spectrometer (Thermo Fisher Scientific, MA, USA) coupled to an EASY-nanoLC 1200 system (Thermo Fisher Scientific, MA, USA). Subsequently, 2 μL peptides of the samples, as well as 1 μg peptide, was loaded and data was pre-collected in DIA mode.

Raw Data of DIA were processed and analyzed by SpectroDive 10.4 (Biognosys AG, Switzerland) with default settings to generate a spectral library. SpectroDive was set up to search the database of Chlorocebus sabaeus which was downloaded from the UniProt database. Library were used to select proteotypic or protein group specific peptides and develop PRM assays by SpectroDive.

Raw files of the targeting runs were analyzed in SpectroDive 10.4 with the default settings. Under the defaults, Q-value cutoff on precursor was applied 1%. The average of filtered peptides was used to calculate the protein quantities.

**Proteomic bioinformatic analysis**

The identified protein groups were filtered and removed duplicate values before further data processing. Protein quantification was calculated by protein raw intensity. Proteins according to the criteria that the fold change > 1.5 and p value < 0.05 were considered differentially expressed proteins. The annotation, gene ontology enrichment, pathway enrichment and bioinformatic analysis were done by OmicShare (https://www.omicshare.com/) and KEGG (https://www.kegg.jp/). The heatmap and PCA analysis were launched by complexheatmap and factoextra package (downloaded from http://bioconductor.org/), respectively.

**Extended Data Figures**

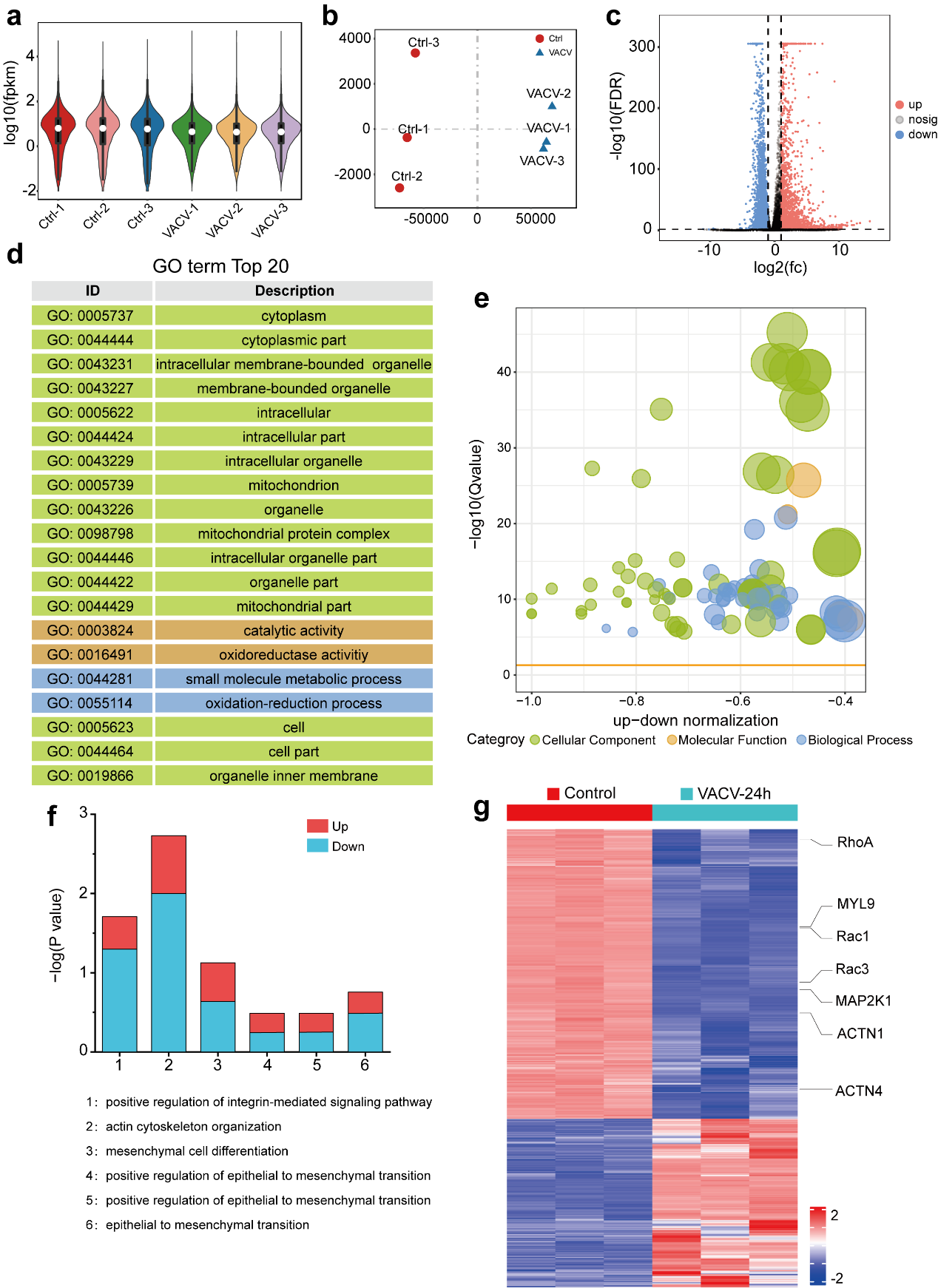

**Extended Data Fig. 1 Control and VACV-infected Vero cells presented transcriptomic differences by RNA-Seq analysis. a,** The violin plot of gene expression visualization of control and VACV-infected Vero cells. The white spot represents the medium intensity and the black line represents the section between maximum and minimum limits. The calculation of gene expression uses the Fragments Per Kilobase of transcript per Million mapped reads (FPKM). FPKM=106C/(NL/103). **b,** The Principal Component Analysis (PCA) plot of control and VACV-infected Vero cells. **c,** The volcano plot of Differentially Expressed Genes (DEGs; fold change > 2 and FDR < 0.05) from control and VACV-infected Vero cells. **d,** The top 20 terms of GO enrichment of DEGs. **e,** The bubble diagram of three main categories of GO enrichment, including biological process, molecular function, and cellular component, by up- and down-regulated genes from control and VACV-infected Vero cells. **f,** GO terms of DEGs from control and VACV-infected Vero cells. The plot shows the p value of overrepresented GO terms associated with EMT. **g**, Heatmap of DEGs associated with RhoA-ROCK signaling pathway and regulation of actin cytoskeleton. The associated DEGs are highlighted. Each group has three biological replicates.

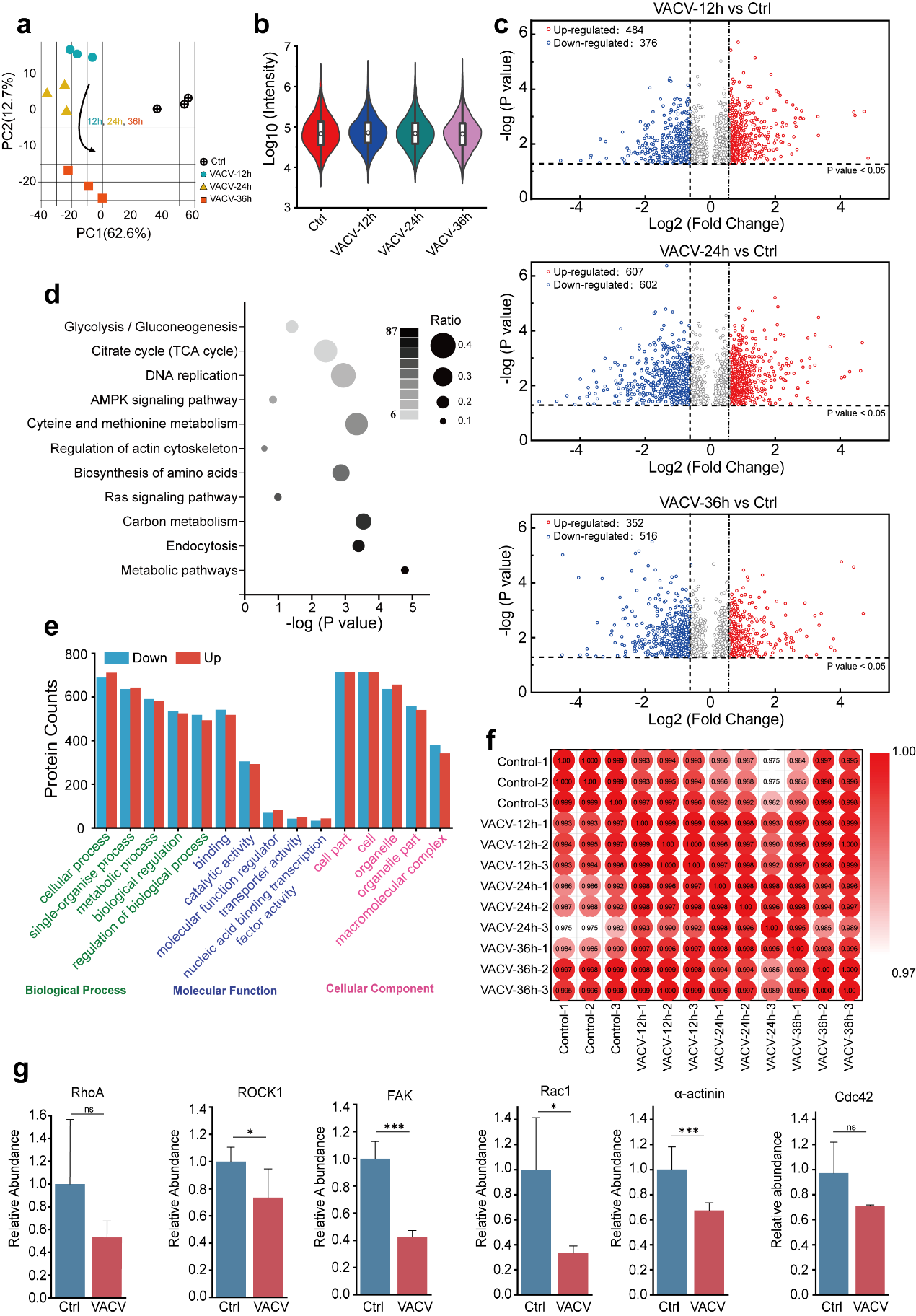

**Extended Data Fig. 2 Differentially expressed proteins in control and VACV-infected Vero cells during various infection time periods by proteomic analysis. a,** Proteomics Principle Component Analysis (PCA) plot of control and VACV-infected Vero cells during different infection time periods. **b,** The violin plot of relative protein intensity of control and VACV-infected Vero cells (12h, 24h, 36h). **c,** The Volcano plot of Differentially Expressed Proteins (DEPs; fold change≧1.5, p value≦0.05) from control and VACV-infected Vero cells (12h, 24h, 36h), respectively. **d,** The bubble diagram of KEGG pathway enrichment of DEPs. The circle size and color shade represent enriched protein ratio and counts in each pathway (from 0.1 to 0.4, small to big), respectively. **e,** Gene Ontology terms, including biological process, molecular function and cellular component enriched DEPs (red represents up-regulated proteins and blue represents down-regulated proteins) of control and VACV-infected Vero cells (12 h, 24 h, 36 h). **f,** The sample correlation plot of control and VACV-infected Vero cells. The color represents the correlation between sample groups (from white to red, weak to strong). **g,** The relative intensity of RhoA signaling pathway-associated proteins in three biological replicates. Data are mean ± SEM. *p<0.05, **p<0.01, ***p<0.001 (unpaired t-test).

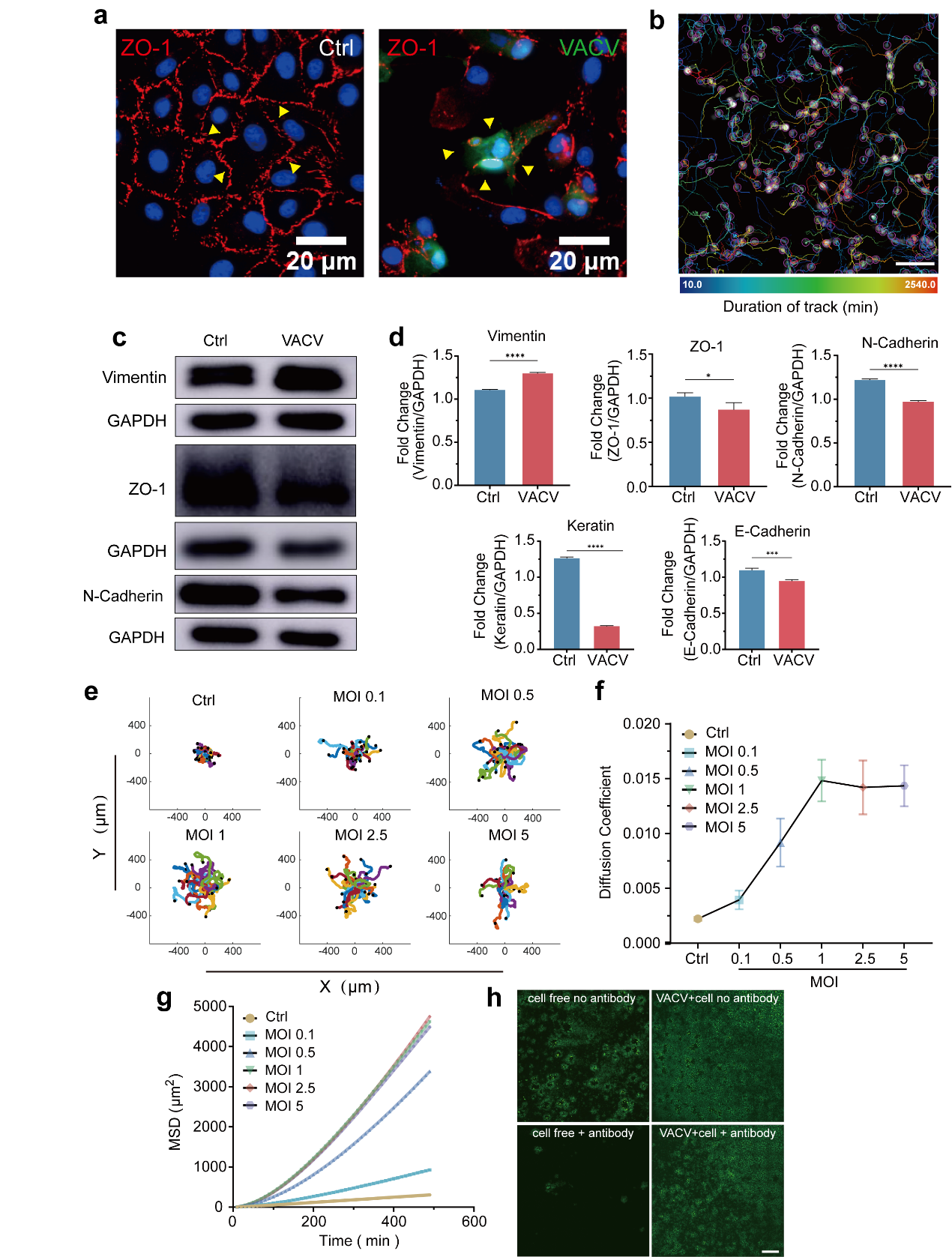

**Extended Data Fig. 3 EMT-like transformation and cell motility after VACV infection. a,** Representative images of the location of ZO-1 (red) in control or VACV-infected Vero cells. **b,** Representative image showing cell trajectory tracked by nucleus location in approximately 43h. Different color means the different duration of the track. Scale bar (h) represents 50 μm. **c, d,** Immunoblotting (c) and quantitative analysis (d) of Vimentin, ZO-1, N-Cadherin, Keratin, and E-Cadherin in Vero cells at 24 h.p.i. (n=3). **e-g,** Normalized trajectory (e), diffusion coefficient (f), and MSD (g) of the migrating Vero cells with different MOI of VACV. **h,** Representative images of VACV-infected Vero cells expressing EGFP in transwell-based virus spread assay. Scale bar (h) represents 1 mm. Data are mean ± SEM. *p<0.05, **p<0.01, ***p<0.001, ****p<0.0001 (unpaired t-test).

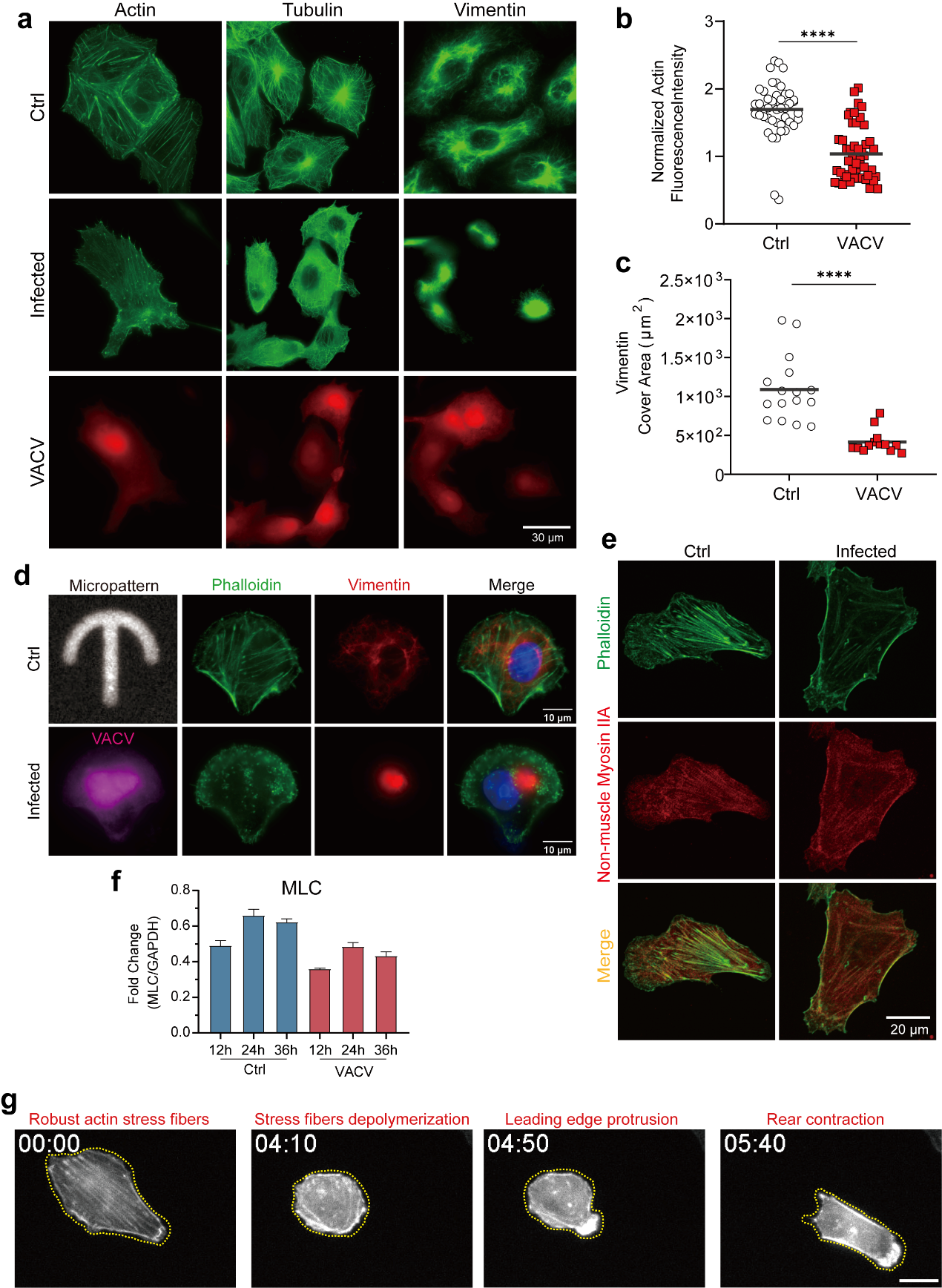

**Extended Data Fig. 4 Actin cytoskeleton is important for VICM. a,** The representative fluorescent images (actin left column; tubulin middle column; vimentin right column) of Vero cells with or without VACV infection. **b,c,** Quantitative analysis of actin fluorescence intensity per cell (b) and vimentin cover area (c) in control and VACV-infected cells. **d,** Representative fluorescent images (actin green; vimentin red; micropattern gray) of control or VACV-infected Vero cells on umbelliform micropatterned bottom. **e,** Representative fluorescent images (actin green; non-muscle myosin IIA red) of Vero cells with or without VACV infection. **f,** Immunoblotting of MLC in control and VACV-infected Vero cells. **g,** Representative time lapse images of a VACV-infected Vero cell expressing LifeAct-mRuby. Scale bar represents 20 μm. Data are mean ± SEM. *p<0.05, **p<0.01, ***p<0.001, ****p<0.0001 (unpaired t-test).

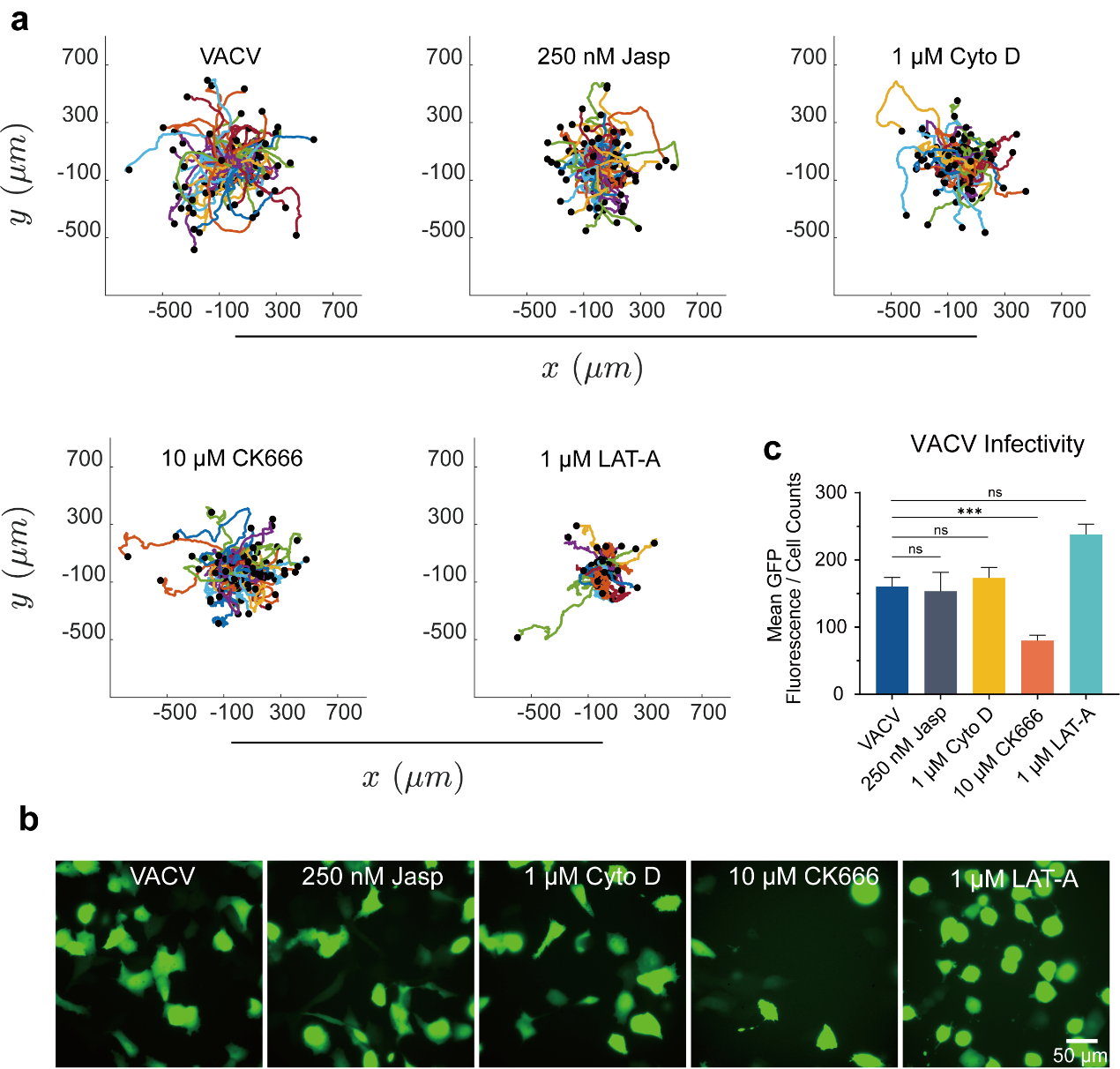

**Extended Data Fig. 5 Actin assemble-related inhibitors affect VICM. a,** The trajectory of VACV-infected Vero cells with actin cytoskeleton inhibitors treatment. **b,c,** The representative fluorescent images (b) and fluorescent quantitative analysis (c) of VACV-infected Vero cells (green) with actin cytoskeleton inhibitors (250 nM Jasp, 1 μM Cyto D, 10 μM CK666, and 1 μM LAT-A) treatment. Data are mean ± SEM. *p<0.05, **p<0.01, ***p<0.001, ****p<0.0001 (unpaired t-test).

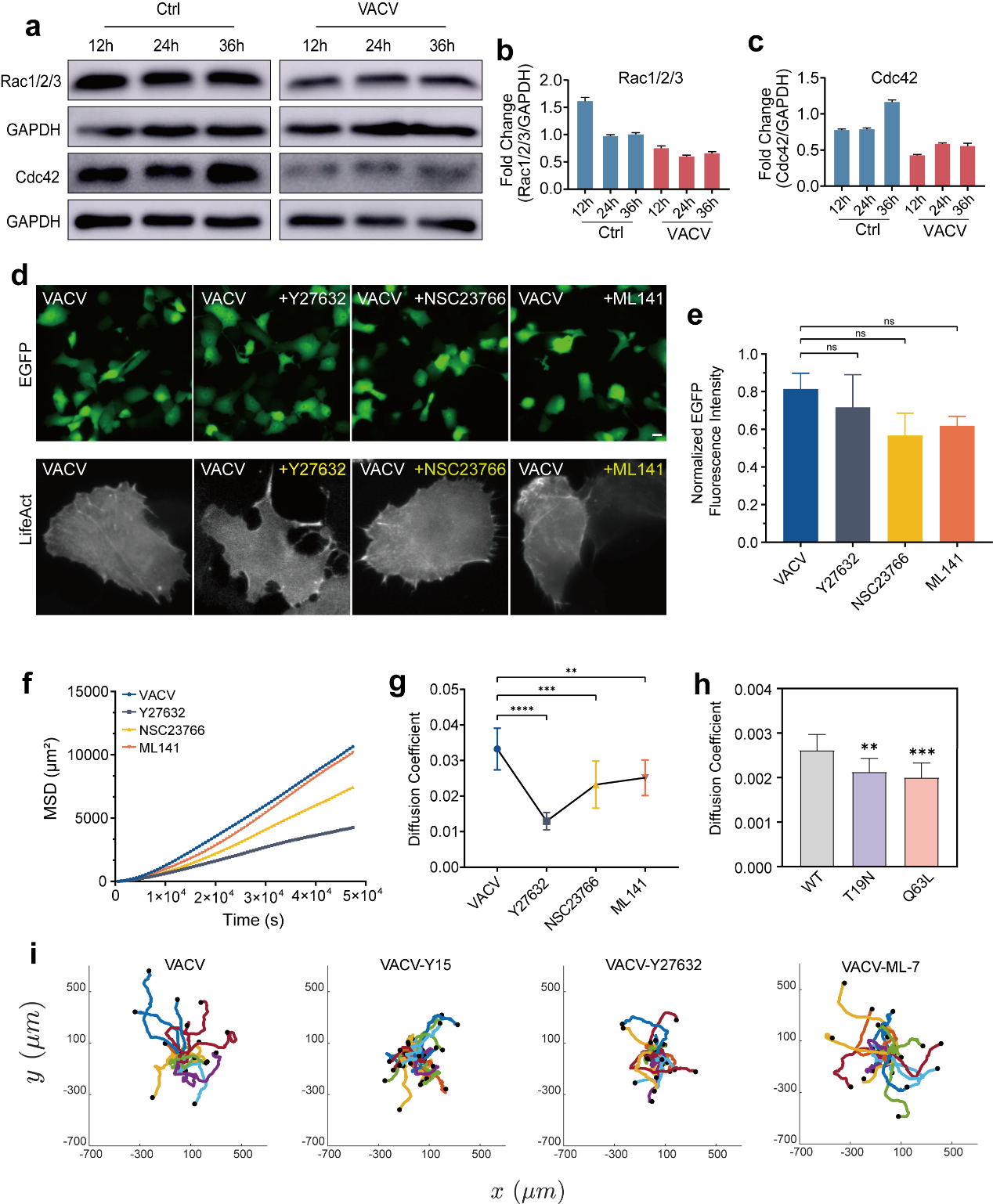

**Extended Data Fig. 6 RhoA and FAK regulate VICM. a-c,** Immunoblotting (a) and quantitative analysis (b, c) of Rac1/2/3 and Cdc42 in Vero cells at 12, 24, and 36 h.p.i. (n=3). **d,** Representative fluorescent images of VACV-infected (top: EGFP) Vero cells expressing LifeAct-mRuby (bottom) with GTPase Inhibitors. Scale bars represent 20 μm. **e,** The fluorescent quantitative analysis of VACV-infected Vero cells (green) with GTPase Inhibitors. **f,g,** MSD (f) and diffusion coefficient (g) of VACV-infected Vero cells with GTPase Inhibitors (80 μM Y27632, 250 μM NSC23766, and 500 nM ML141). **h,** Diffusion coefficient of VACV-infected Vero cells expressing RhoA T19N or Q63L. **i,** The trajectory of VACV-infected Vero cells with inhibitors of FAK (Y15), ROCK (Y27632), and ML-7 (MLCK). Data are mean ± SEM. *p<0.05, **p<0.01, ***p<0.001 (unpaired t-test).

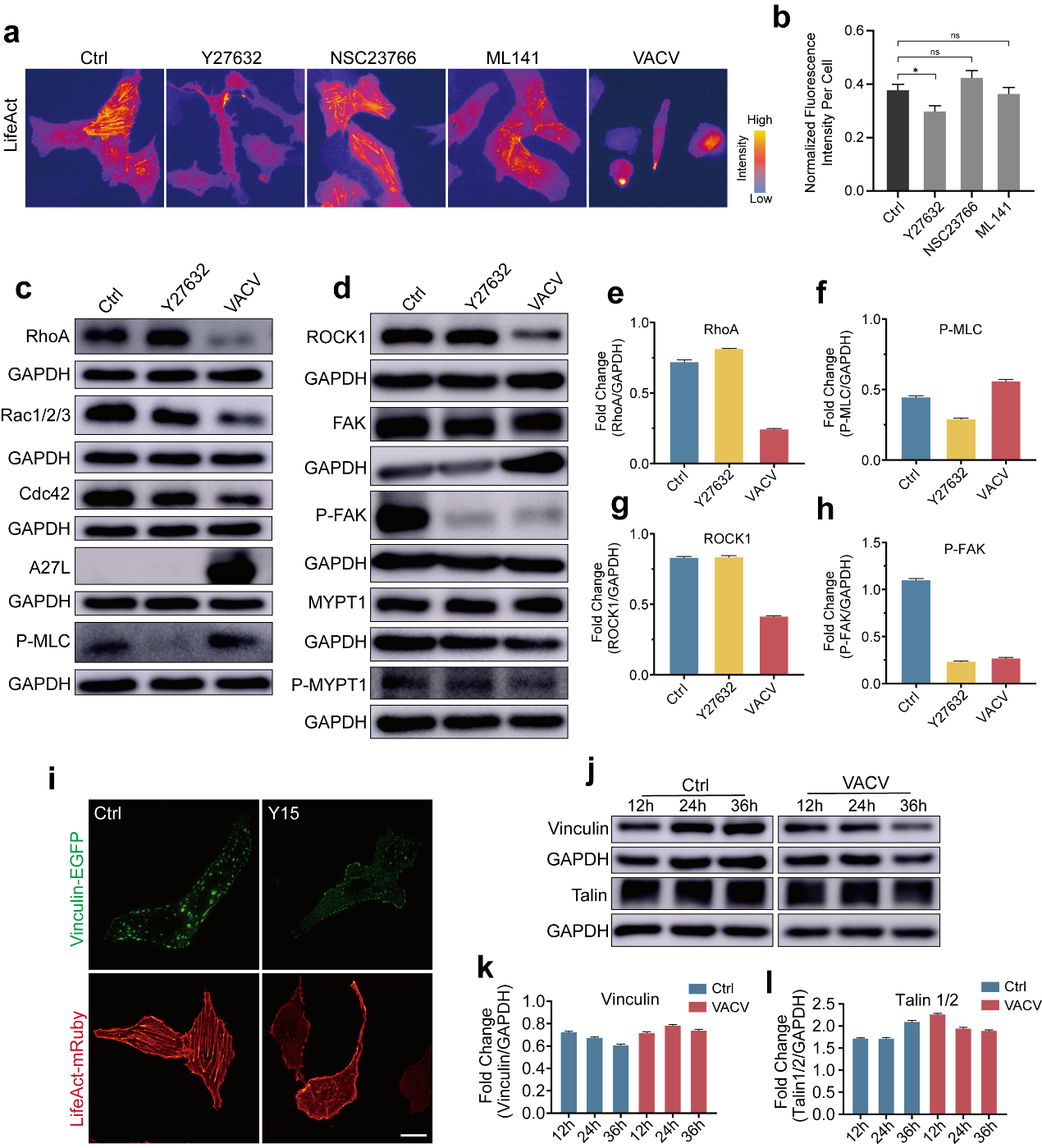

**Extended Data Fig. 7 RhoA signaling is responsible for actin cytoskeleton dynamics during VICM. a,b,** The representative fluorescent images (a) and fluorescent quantitative analysis (b) of Vero-LifeAct-mRuby cells with GTPase Inhibitors or VACV infection. Yellow arrows mean the protrusion was generating around the cells. Scale bar represents 20 μm. **c-h,** Immunoblotting (c,d) and quantitative analysis (e-h) of RhoA, Rac1/2/3, Cdc42, ROCK1, FAK, p-FAK, MYPT1, p-MYPT1 and p-MLC in Vero cells with Y27632 or VACV treatment (n=3). **i,** Representative fluorescent images of Vero cells expressing vinculin-EGFP or LifeAct-mRuby after Y15 treatment. Scale bar represents 20 μm. **j-l,** Immunoblotting (j) and quantitative analysis (k,l) of vinculin and talin in VACV-Vero cells at 12, 24 and 36 h.p.i. (n=3). Data are mean ± SEM. *p<0.05, (unpaired t-test).

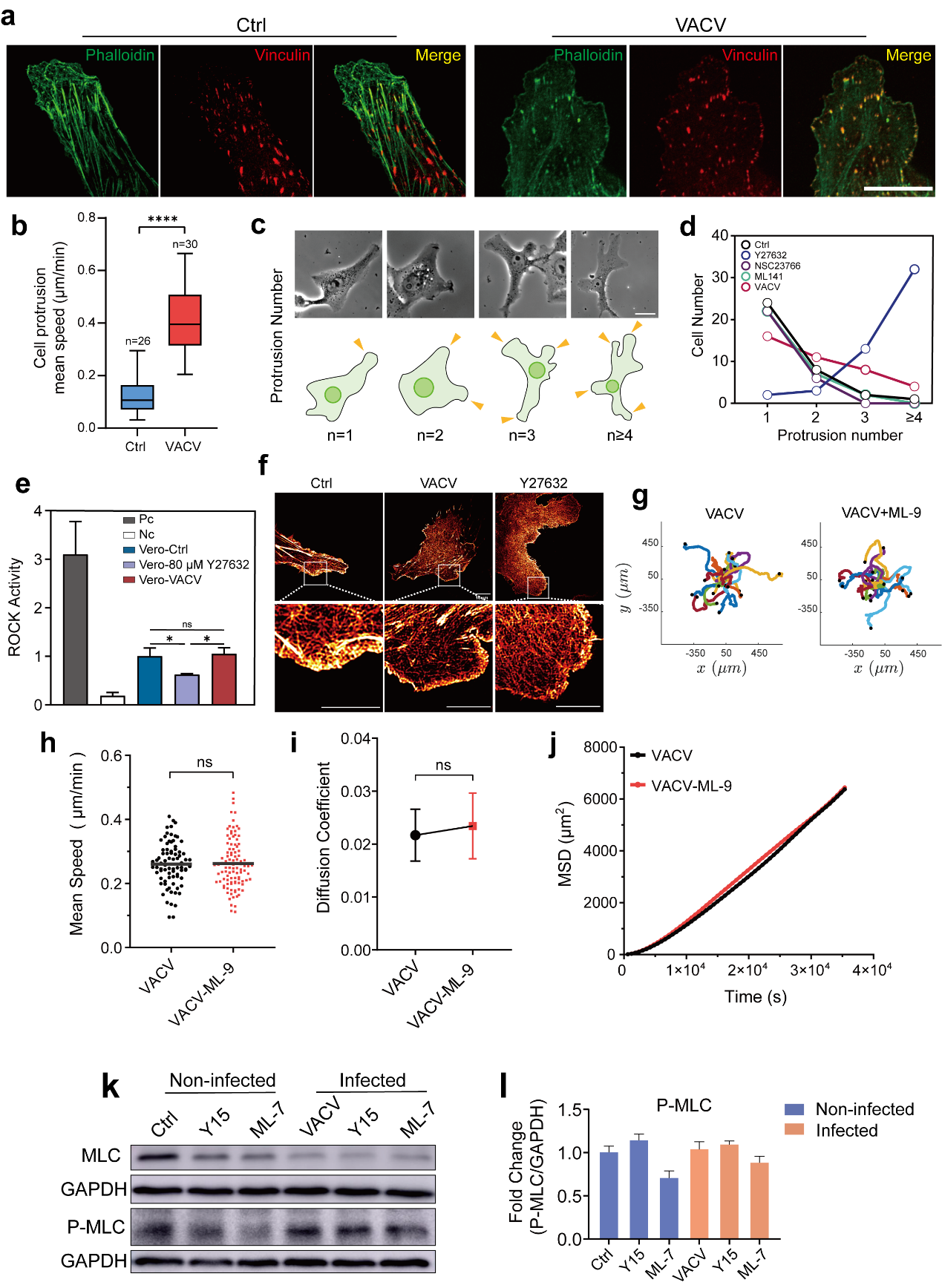

**Extended Data Fig. 8 VACV-regulated RhoA signaling is responsible for protrusion formation and cell contractility during VICM. a,** Representative fluorescent images (actin green; vinculin red) of control or VACV-infected Vero cells around the protrusion. **b,** The quantitative analysis of extended protrusion in non-infected and VACV infected cells. **c,** Representative images and schematic diagram of cells with different protrusion number. Yellow arrows indicate the protrusions in the cells. Scale bar represents 20 μm. **d,** The relationship between cell number and protrusion number after inhibitors or VACV treatment. **e,** Quantitative analysis of ROCK activity in Vero cells with different treatment (Pc: positive control; Nc: negative control). **f,** Representative super-resolution images of Vero-LifeAct-mRuby cells under VACV or Y27632 treatment. Scale bars represent 5μm. **g-j,** The trajectory (f), mean speed (g), diffusion coefficient (h), and MSD (i) of VACV-infected Vero cells with ML-9 treatment. **k,** Immunoblotting of MLC and p-MLC in control or VACV-infected Vero cells with inhibitors of Y15 or ML-7 (n=3). **l,** The quantitative analysis of p-MLC in control or VACV-infected Vero cells with inhibitors of Y15 or ML-7. Data are mean ± SEM. *p<0.05, **p<0.01, ***p<0.001, ****p<0.0001 (unpaired t-test).

**Supplementary Video Legends**

**Supplementary Video 1.** Representative recording of the Vero cells expressing LifeAct-mRuby (black) without (left) or with VACV infection (right), related to Fig. 2f. Scale bars represent 30 μm.

**Supplementary Video 2.** Representative recording of the actin dynamic in a VACV-infected Vero cell expressing LifeAct-mRuby (gray), related to Extended Data Fig. 4g. Scale bar represents 30 μm.

**Supplementary Video 3.** Spinning disk confocal microscopy recording of the actin dynamic in a Vero cell expressing LifeAct-mRuby (royal), related to Fig. 2i. Scale bar represents 10 μm.

**Supplementary Video 4.** Spinning disk confocal microscopy recording of the actin dynamic in a VACV-infected Vero cell expressing LifeAct-mRuby (royal), related to Fig. 2k. Scale bar represents 10 μm.

**Supplementary Video 5.** Representative recording of the VACV-infected Vero cells expressing LifeAct-mRuby (gray) with different inhibitors treatment (250 nM Jasplakinolide, 1 μM Cytochalasin D, 1 μM Latrunculin A, 10 μM CK666 added at 8 h.p.i.), related to Fig. 2n. Scale bars represent 30 μm.

**Supplementary Video 6.** Spinning disk confocal microscopy recording of the dynamic of robust actin stress fibers depolymerization and protrusion formation in VACV-infected Vero cells expressing LifeAct-mRuby (gray), related to Fig. 4a and 5a. Scale bar represents 20 μm.

**Supplementary Video 7.** Spinning disk confocal microscopy recording of actin stress fibers in Vero cells expressing LifeAct-mRuby（black）without (left) or with VACV infection (right), related to Fig. 4e. Scale bars represent 10 μm.

**Supplementary Video 8.** Spinning disk confocal microscopy recording of the dynamic of protrusion in a non-infected Vero cell expressing LifeAct-mRuby (red), related to Fig. 5b. Scale bar represents 20 μm.

**Supplementary Video 9.** Spinning disk confocal microscopy recording of the dynamic of protrusion in a VACV-infected Vero cell expressing LifeAct-mRuby (red), related to Fig. 5c. Scale bar represents 20 μm.

**Supplementary Video 10.** TIRF microscopy recording of the dynamic of focal adhesion in the non-infected Vero cells expressing vinculin-EGFP (gray), related to Fig. 5d. Scale bar represents 20 μm.

**Supplementary Video 11.** TIRF microscopy recording of the dynamic of focal adhesion in VACV-infected Vero cells expressing vinculin-EGFP (gray), related to Fig. 5d. Scale bar represents 20 μm.

**Supplementary Video 12.** Spinning disk confocal microscopy recording of the dynamic of protrusion in 80 μM Y27632 treated Vero cells expressing LifeAct-mRuby (gray), related to Fig. 5e. Scale bar represents 20 μm.

**Supplementary Video 13.** Spinning disk confocal microscopy recording of the dynamic of protrusion in 10 μM Y27632 treated Vero cells expressing LifeAct-mRuby (gray), related to Fig. 5e. Scale bar represents 20 μm.

**Supplementary Video 14.** Spinning disk confocal microscopy recording of the dynamic of protrusion in 10 μM Y15 treated Vero cells expressing LifeAct-mRuby (gray), related to Fig. 5e. Scale bar represents 20 μm.

**Supplementary Video 15.** Spinning disk confocal microscopy recording of the dynamic of the rear in Vero cells expressing LifeAct-mRuby (black) without (left) or with VACV infection (right), related to Fig. 6c. Scale bars represent 10 μm.

**Supplementary Video 16.** Spinning disk confocal microscopy recording of 80 μM Y27632 treated Vero cells expressing LifeAct-mRuby (black), related to Fig. 6e. Scale bar represents 20 μm.
